# Cell Assembly Dynamics of Sparsely-connected Inhibitory Networks: a Simple Model for the Collective Activity of Striatal Projection Neurons

**DOI:** 10.1101/036608

**Authors:** David Angulo-Garcia, Joshua D. Berke, Alessandro Torcini

## Abstract

Striatal projection neurons form a sparsely-connected inhibitory network, and this arrangement may be essential for the appropriate temporal organization of behavior. Here we show that a simplified, sparse inhibitory network of Leaky-Integrate-and-Fire neurons can reproduce some key features of striatal population activity, as observed in brain slices. In particular we develop a new metric to determine the conditions under which sparse inhibitory networks form anti-correlated cell assemblies with time-varying activity of individual cells. We find that under these conditions the network displays an input-specific sequence of cell assembly switching, that effectively discriminates similar inputs. Our results support the proposal that GABAergic connections between striatal projection neurons allow stimulus-selective, temporally-extended sequential activation of cell assemblies. Furthermore, we help to show how altered intrastriatal GABAergic signaling may produce aberrant network-level information processing in disorders such as Parkinson’s and Huntington’s diseases.

**Author Summary:** Neuronal networks that are loosely coupled by inhibitory connections can exhibit potentially useful properties. These include the ability to produce slowly-changing activity patterns, that could be important for organizing actions and thoughts over time. The striatum is a major brain structure that is critical for appropriately timing behavior to receive rewards. Striatal projection neurons have loose inhibitory interconnections, and here we show that even a highly simplified model of this striatal network is capable of producing slowly-changing activity sequences. We examine some key parameters important for producing these dynamics, and help explain how changes in striatal connectivity may contribute to serious human disorders including Parkinson’s and Huntington’s diseases.

## Introduction

The basal ganglia are critical brain structures for behavioral control, whose organization has been highly conserved during vertebrate evolution [1]. Altered activity of the basal ganglia underlies a wide range of human neurological and psychiatric disorders, but the specific computations normally performed by these circuits remain elusive. The largest component of the basal ganglia is the striatum, which appears to have a key role in adaptive decision-making based on reinforcement history [2], and in behavioral timing on scales from tenths of seconds to tens of seconds [3].

The great majority (> 90%) of striatal neurons are GABAergic medium spiny neurons (MSNs), which project to other basal ganglia structures but also make local collateral connections within striatum [4]. These local connections were proposed in early theories to achieve action selection through strong winner-take-all lateral inhibition [5, 6], but this idea fell out of favor once it became clear that MSN connections are actually sparse (nearby connection probabilities ≃ 10 – 25% [7, 8]), unidirectional and relatively weak [9, 10]. Nonetheless, striatal networks are intrinsically capable of generating sequential patterns of cell activation, even in brain slice preparations without time-varying external inputs [11, 12]. Following previous experimental evidence that collateral inhibition can help organize MSN firing [13], an important recent set of modeling studies argued that the sparse connections between MSNs, though individually weak, can collectively mediate sequential switching between cell assemblies [14, 15]. It was further hypothesized that these connections may even be optimally configured for this purpose [16]. This proposal is of high potential significance, since sequential dynamics may be central to the striatum’s functional role in the organization and timing of behavioral output [17, 18].

In their work [14, 15, 16], Ponzi and Wickens used conductance-based model neurons (with persistent *Na*^+^ and *K*^002B;^ currents [19]), in proximity to a bifurcation from a stable fixed point to a tonic firing regime. We show here that networks based on simpler leaky integrate-and-fire (LIF) neurons can also exhibit sequences of cell assembly activation. This simpler model, together with a novel measure of structured bursting, allows us to more clearly identify the critical parameters needed to observe dynamics resembling that of the striatal MSN network. Among other results, we show that the duration of GABAergic post-synaptic currents is crucial for the network’s ability to discriminate different input patterns. GABAergic currents are abnormally brief in mouse models of Huntington’s Disease (HD) [20], and we demonstrate how this may produce the altered neural activity dynamics reported for symptomatic HD mice [21]. Finally, dopamine loss weakens MSN-MSN interactions [22, 23], and our results help illuminate the origins of aberrant synchronization patterns in Parkinson’s Disease (PD).

## Results

### Measuring cell assembly dynamics

The model is composed of N leaky integrate-and-fire (LIF) inhibitory neurons [24, 25], with each neuron receiving inputs from a randomly selected 5% of the other neurons (i.e. a directed ErdÖs-Renyi graph with constant in-degree *K = pN*, where *p* = 5%) [26]. The inhibitory post-synaptic potentials (PSPs) are modeled as *α*-functions characterized by a decay time *τ_α_* and a peak amplitude *A_PSP_*. In addition, each neuron *i* is subject to an excitatory input current mimicking the cortical and thalamic inputs received by the striatal network. In order to obtain firing periods of any duration the excitatory drives are tuned to drive the neurons in proximity of the supercritical bifurcation between the quiescent and the firing state, similarly to [14]. Furthermore, our model is integrated exactly between a spike emission and the successive one by rewriting the time evolution of the network as an event-driven map [27] (for more details see Methods).

**Figure 1.**
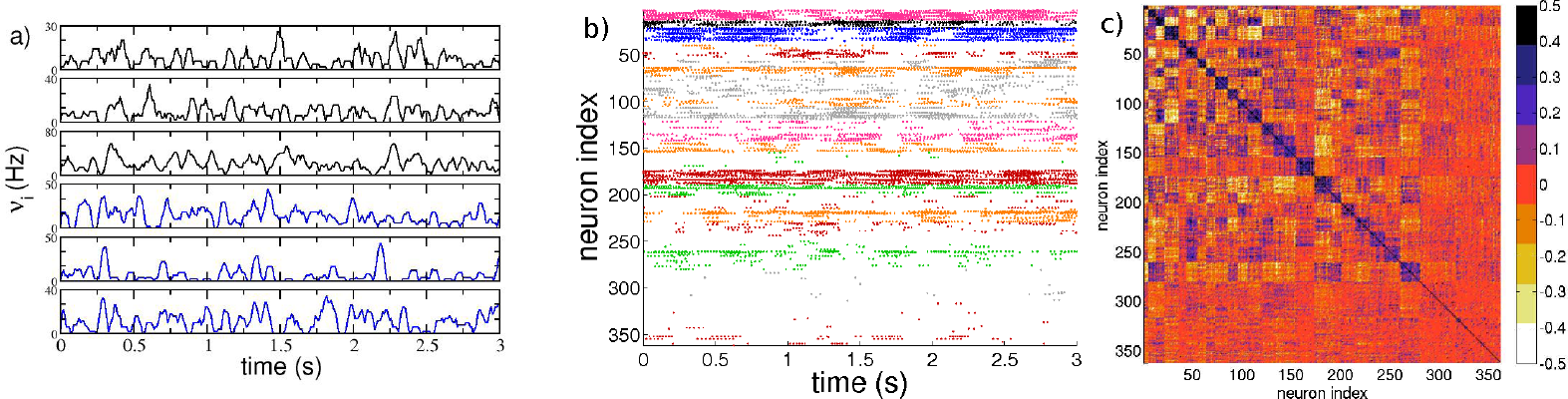
Cell activity characterization. a) Firing rates *ν_i_* of 6 selected neurons belonging to two anti-correlated assemblies, the color identifies the assembly and the colors correspond to the one used in b) for the different clusters; b) raster plot activity, the firing times are colored according to the assembly the neurons belong to; c) cross-correlation matrix C(*υ_i_, υ_j_*) of the firing rates. The neurons in panel b) and c) are clustered according to the correlation of their firing rates by employing the *k-means* algorithm; the clusters are ordered in terms of their average correlation (inside each cluster) from the highest to the lowest one (for more details see Methods). The firing rates are calculated over overlapping time windows of duration 1 s, the origins of successive windows are shifted by 50 ms. The system is evolved during 10^7^ spikes, after discarding an initial transient of 10^5^ spike events. Other parameters used in the simulation: 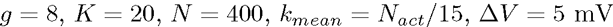 and *τ_α_* = 20 ms. The number of active neurons is 370, corresponding to *n*\* ≃ 93 %.

Since we will compare most of our findings with the results reported in a previous series of papers by Ponzi and Wickens (PW) [14, 15, 16] it is important to stress the similarities and differences between the two models. The model employed by PW is a two dimensional conductance-based model with a potassium and a sodium channel [19], our model is simply a current based LIF model. The parameters of the PW model are chosen so that the cell is in proximity of a saddle-node on invariant circle (SNIC) bifurcation to guarantee a smooth decrease of the firing period when passing from the quiescent to the supra-threshold regime, without a sudden jump in the firing rate. Similarly, in our simulations the parameters of the LIF model are chosen to be in proximity of the bifurcation from silent regime to tonic firing. In the PW model the PSPs are assumed to be exponentially decaying, whereas we considered α-functions.

In particular, we are interested in determining model parameters for which uniformly distributed inputs 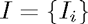, where 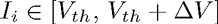, produce cell assembly-like sequential patterns in the network. The main aspects of the desired activity can be summarized as follows: (i) single neurons should exhibit large variability in firing rates (*CV* > 1); (ii) the dynamical evolution of neuronal assemblies should reveal strong correlation within an assembly and anti-correlation with neurons out of the assembly. As suggested by many authors [9, 28] the dynamics of MSNs cannot be explained in terms of a *winners take all* (WTA) mechanism, which would imply a small number of highly firing neurons, while the remaining would be silenced. Therefore we searched for a regime in which a substantial fraction of neurons are actively involved in the network dynamics. This represents a third criterion (iii) to be fulfilled to obtain a striatum-like dynamical evolution.

Fig. 1 shows an example of such dynamics for the LIF model, with three pertinent similarities to previously observed dynamics of real MSN networks [11]. Firstly, cells are organized into correlated groups, and these groups are mutually anticorrelated (as evident from the cross-correlation matrix of the firing rates reported in Fig. 1 (c)). Secondly, individual cells within groups show irregular firing as shown in Fig. 1 (a). This aspect is reflected in a coefficient of variation (*CV*) of the inter-spike-intervals (ISIs) definitely greater than one (see the black curve in Fig. 3 (b)) as observed experimentally for the dynamics of rat striatum *in-vitro* [7, 9]. Thirdly, the raster plot reported in Fig. 1 (b) reveals that a large fraction of neurons (namely, ≃ 93 %) is active.

### A novel metric for the structured cell assembly activity

The properties (i),(ii), and (iii), characterizing MSN network activity, can be quantified in terms of a single scalar metric *Q*_0_, as follows:

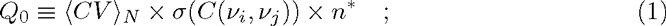

where ⟨•⟩*N* denotes average over the ensemble of *N* neurons, *n*\* = *N_act_/N* is the fraction of active neurons *N_act_* out of the total number, *C*(*υ_i_, υ_j_*) is the *N χ N* zero-lag cross-correlation matrix between all the pairs of single neuron firing rates (*υ_i_, υ_j_*), and *σ*(•) is the standard deviation of this matrix (for details see Methods). We expected that suitable parameter values for our model could be selected by maximizing *Q*_0_.

Our metric is inspired by a metric introduced to identify the level of cluster synchronization and organization for a more detailed striatal microcircuit model [29]. However, that metric is based on the similarity among the point-process spike trains emitted by the various neurons, whereas *Q*_0_ uses correlations between firing rate time-courses. Furthermore, *Q*_0_ takes also in account the variability of the firing rates, by including the average *CV* in Eq. (1), an aspect of the MSN dynamics omitted by the metric employed in [29]. Within biologically meaningful ranges, we find values of the parameters controlling lateral inhibition (namely, the synaptic strength *g* and the the post-synaptic potential duration *τ_α_*) that maximize *Q*_0_. As we show below, the chosen parameters not only produce MSN-like network dynamics but also optimize the network’s computational capabilities, in the sense of producing a consistent, temporally-structured response to a given pattern of inputs while discriminating between very similar inputs.

### The role of lateral inhibition

In this sub-section we examine how network dynamics are affected by the strength of inhibitory connections (Fig. 2). When these lateral connections are very weak (parameter *g* close to zero), the dominant input to each neuron is the constant excitation. As a result, most individual neurons are active (fraction of active neurons, *n*\*, is close to 1) and firing regularly (*CV* close to zero). As lateral inhibition is made stronger, some neurons begin to slow down or even stop firing, and *n*\* declines towards a minimum fraction of ≃ 50% (at *g = g_min_*). As noted by Ponzi and Wickens [16], this is due to a winner-take-all (WTA) mechanism: faster-firing neurons depress or silence the neurons to which they are connected. This is evident from the distribution 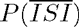 of the average interspike intervals 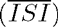, which is peaked at low firing periods, and from the distribution of the *CV* exhibiting a single large peak at *CV* ≃ 0 (as shown in the insets of Figs. S2 (a,b) and (d,e)).

As soon as *g* approaches *g_min_*, the neuronal activity is no longer due only to the *winners*, but also the *losers* begin to play a role. The *winners* are characterized by an effective input *W_i_* which is on average supra-threshold, while their firing activity is driven by the mean current: *winners* are *mean-driven* [30]. On the other hand, *losers* are on average below-threshold, and their firing is due to current fluctuations: *losers* are *fluctuation-driven* [30]. For more details see Figs. S2 (c) and (f)). This is reflected in the corresponding distribution 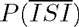 (Fig. 2(b), red curve). The *winners* have very short 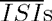 (i.e. high firing rates), while the *losers* are responsible for the long tail of the distribution extending up to 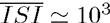. In the distribution of the coefficients of variation (Fig. 2(b) inset, red curve) the *winners* generate a peak of very low *CV* (i.e. highly-regular firing), suggesting that they are not strongly influenced by the other neurons in the network and therefore have an effective input on average supra-threshold. By contrast the *losers* are associated with a smaller peak at *CV* ≃ 1, confirming that their firing is due to large fluctuations in the currents.

Counterintuitively however, further increases in lateral inhibition strength result in increased neuronal participation, with *n*\* progressively returning towards ≃ 1. The same effect was previously reported by Ponzi and Wickens [16] for a different, more complex, model. When the number of active neurons returns almost to 100%, i.e. for sufficiently large coupling *g > g_min_*, most of the neurons appear to be below threshold, as revealed by the distribution of the effective inputs *W_i_* reported in Figs. S2 (c) and (f). Therefore in this case the network firing is essentially fluctuation-driven, and the 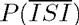 distribution is now characterized by a broader distribution and by the absence of the peak at short 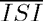 (as shown in Fig. 2 (b), blue line; see also Figs. S2(a) and (d)). Furthermore the single neuron dynamics is definitely bursting, as shown by the *CV* distribution now centered around *CV* ≃ 2 (inset of Fig. 2 (b), blue line; see also Figs. S2(b) and (e)).

The transition between these two dynamical regimes, occurring at *g = g_min_*, is due to a passage from a state where some *winner* neurons were mean-driven and able to depress all the other neurons, to a state at *g >> g_min_* where almost all neurons are fluctuation-driven and contribute to the network activity. The transition occurs because at *g < g_min_* the fluctuations in the effective input currents *W_i_* are small and insufficient to drive the *losers* towards the firing threshold (as shown in the insets of Fig. S2 (c) and (f)). At *g ≃ g_min_* the amplitude of the fluctuations becomes sufficient for some *losers* to cross the firing threshold and contribute to the number of active neurons. This will also reduce the *winners’* activity. For *g >> g_min_* the fluctuations of *W_i_* are sufficient to lead almost all *losers* to fire and no clear distinction between *losers* and *winners* remains.

The reported results explain why the variability *σ*(*C*) of the cross-correlation matrix has a non monotonic behaviour with *g* (as shown in the middle panel in Fig. 2(a)). At low coupling *σ*(*C*) is almost zero, since the single neuron dynamics are essentially independent one from another, while the increase of the coupling leads to an abrupt rise of *σ*(*C*). This growth is clearly associated with the inhibitory action which splits the neurons into correlated and anti-correlated groups. The variability of the cross-correlation matrix achieves a maximum value for coupling slightly larger than *g_min_*, where fluctuations in the effective currents begin to affect the network dynamics. At larger coupling *σ*(*C*) begins to decay towards a finite non zero value. These results confirm that the most interesting region to examine is the one with coupling *g > g_min_*, as already suggested in [16].

The observed behaviour of *CV, n*\* and *σ*(*C*) suggests that we should expect a maximum in *Q*_0_ at some intermediate coupling *g > g_min_*, as indeed we have for both studied cases as shown in Fig. 2 (c) and (d). The initial increase in *Q*_0_ is due to the increase in *CV* and *n*\*, while the final decrease, following the occurrence of the maximum, is essentially driven by the decrease of *σ*(*C*). For larger Δ*V* the neurons tend to fire regularly across a wider range of small *g* values (see Fig. 2 (d)), indicating that due to their higher firing rates a larger synaptic inhibition is required to influence their dynamics. On the other hand, their bursting activity observable at large *g* is more irregular (see the upper panel in Fig. 2 (a); dashed line and empty symbols).

To assess whether parameters that maximize *Q*_0_ also allow discrimination between different inputs, we alternated the network back and forth between two distinct input patterns, each presented for a period *T_sw_*. During this stimulation protocol, we evaluated the state transition matrix (STM) and the associated quantity Δ*M_d_*. The STM measures the similarity among the firing rates of the neurons in the network taken at two different times. The metric Δ*M_d_*, based on the STM, has been introduced in [16] to quantify the ability of the network to distinguish between two inputs. In particular, Δ*M_d_* is the difference between the average values of the STM elements associated with the presentation of each of the two stimuli (a detailed description of the procedure is reported in the sub-section *Discriminative and computation capability* and in Methods).

To verify whether the ability of the network to distinguish different stimuli is directly related to the presence of dynamically correlated and anti-correlated cell assemblies, we generated a new metric, *Q_d_*. This metric is defined in the same way as *Q*_0_, except that in Eq. (1) the standard deviation of the correlation matrix is replaced by Δ*M_d_*. As it can be appreciated from Figs. 2(c) and 2(d) the metrics *Q_d_* and *Q*_0_ behave similarly, indicating that indeed *Q*_0_ becomes maximal in the parameter range in which the network is most effectively distinguishing different stimuli. We speculate that the emergence of correlated and anti-correlated assemblies contributes to this discriminative ability.

We note that we observed maximal values of *Q*_0_ for realistic lateral inhibition strengths, as measured from the post-synaptic amplitudes *A_PSP_*. Specifically, *Q*_0_ reaches the maximum at *g* = 4 (*g* = 8) for Δ*V* = 1 mV (Δ*V* = 5 mV) corresponding to *A_PSP_* = 0.368 mV (*A_PSP_* = 0.736 mV), comparable to previously reported experimental results [7, 28, 9] (for more details see Methods).

### The role of the post-synaptic time scale

In brain slice experiments IPSCs/IPSPs between MSNs last 5-20 ms and are mainly mediated by the GABA*a*-receptor [7, 31]. In this sub-section, we will examine the effect of the the post-synaptic time constant *τ_α_*. As *τ_α_* is increased from 2 to 50 ms, the values of of both metrics *Q*_0_ and *Q_d_* progressively increase (Fig. 3(a)), with the largest variation having already occurred by *τ_α_* = 20 ms. To gain more insights on the role of the PSP in shaping the structured dynamical regime, we show for the same network the distribution of the single cell *CV*, for *τ_α_* = {2, 9, 20} ms (Fig. 3(b)). Narrow pulses (*τ_α_* ≃ 2 ms) are associated with a distribution of *CV* values ranging from 0.5 to 1, with a predominant peak at one. By increasing *τ_α_* one observes that the *CV* distributions shift to larger and larger *CV* values. Therefore, one can conclude that at small *τ_α_* the activity is mainly Poissonian, while increasing the duration of the PSPs leads to bursting behaviours, as in experimental measurements of the MSN activity [21]. In particular in [21], the authors showed that bursting activity of MSNs with a distribution *P*(*CV*) centered around *CV* ≃ 2 is typical of awake wild-type mice. To confirm this analysis we estimated the distribution of *CV*_2_: A *CV*_2_ distribution with a peak around zero denotes very regular firing, while a peak around one indicates the presence of long silent periods followed by rapid firing events (i.e. a bursting activity). Finally a flat distribution denotes Poissonian distributed spiking. It is clear from Fig. 3(c) that by increasing *τ_α_* from 2 to 20 ms this leads the system from an almost Poissonian behaviour to bursting, where almost regular firing inside the burst (intra-burst) is followed by a long quiescent period (inter-burst) before starting again.

**Figure 2.**
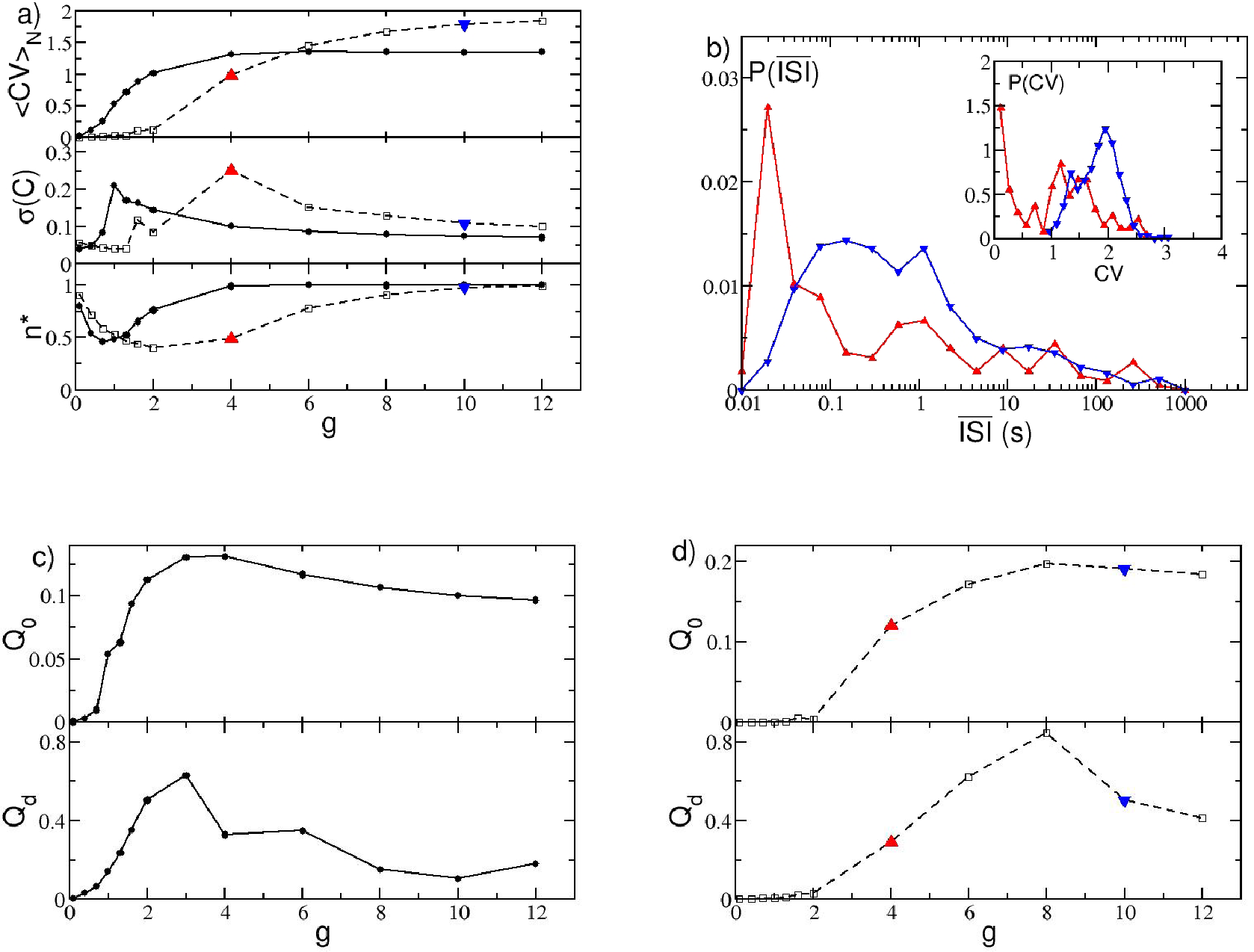
Metrics of structured activity vs lateral inhibition strength. a) Metrics entering in the definition of *Q*_0_ versus the synaptic strength *g*. From top to bottom: Averaged coefficient of variation ⟨*CV*⟩*_N_*, standard deviation of the cross-correlation matrix *σ*(*C*), and the fraction of active neurons *n*\*. The solid (dashed) line refers to the case Δ*V* = 1 mV (Δ*V* = 5 mV). The minimum number of active neurons is achieved at *g = g_min_*, this corresponds to a peak amplitude of the PSP *A_PSP_* = 0.064 mV (*A_PSP_* = 0.184 mV) for Δ*V* = 1 mV (Δ*V* = 5 mV) (for more details see Methods). b) Distributions 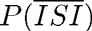 of the average ISI for a fixed Δ*V* = 5 mV and for two different coupling strengths, *g* = 4 (red triangle-up symbol) and *g* = 10 (blue triangle-down). Inset, the distribution *P*(*CV*) of the *CV* of the single neurons for the same two cases. c) *Q*_0_ and *Q_d_*, as defined in Eqs. (1) and (16), versus *g* for Δ*V* = 1 mV. d) Same as c) for Δ*V* = 5 mV. Other parameters as in Fig. 1

**Figure 3.**
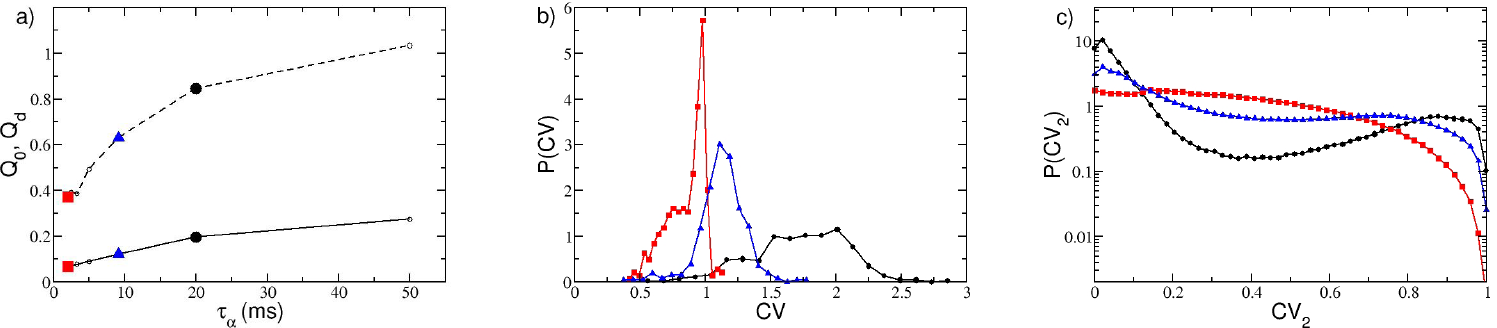
Metrics of structured activity vs post-synaptic time duration. a) Metrics *Q*_0_ (in solid line) and *Q_d_* (dashed) as a function of the pulse time scale for the parameter values {Δ*V, g*} = {5 mV, 8} corresponding to the maximum *Q*_0_ value in Fig. 2(d). Probability distribution functions *P*(*CV*) (*P*(*CV*_2_)) for the coefficient of variation *CV* (local coefficient of variation *CV*_2_) are shown in b) (in c)) for three representative *τ_α_* = {2, 9, 20} ms, displayed by employing the same symbols and colors as indicated in a). For these three cases the average firing rate in the network is ⟨*v*⟩ = {8.81, 7.65, 7.35} Hz ordered for increasing *τ_α_*-values. Other parameters as in Fig. 1

The distinct natures of the distributions of *CV* for short and long pulses raises the question of what mechanism underlies such differences. To answer this question we analyzed the distribution of the ISI of a single cell in the network for two cases: in a cell assembly bursting regime (corresponding to *τ_α_* = 20 ms) and for Poissonian unstructured behavior (corresponding to *τ_α_* = 2 ms). We expect that even the single neurons should have completely different dynamics in these two regimes, since the distributions *P*(*CV*) at *τ_α_* =2 ms and 20 ms are essentially not overlapping, as shown in Fig. 3(b). In order to focus the analysis on the effects due to the synaptic inhibition, we have chosen, in both cases, neurons receiving exactly the same external excitatory drive *I_s_*. Therefore, in absence of any synapses, these two neurons will fire with the same period 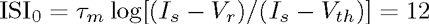 ms, corresponding to a firing rate of 8.33 Hz not far from the average firing rate of the networks (namely, 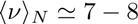 Hz). Thus these neurons can be considered as displaying typical activity in both regimes. As expected, the dynamics of the two neurons is quite different, as evident from the *P*(*ISI*) presented in Fig. 4(a) and (b). In both cases one observes a long tailed exponential decay (with decay rate *v_D_*) of *P*(*ISI*) corresponding to a Poissonian like behaviour. However the decay rate *v_D_* is slower for *τ_α_* = 20 ms with respect to *τ_α_* = 2 ms, namely *v_D_* ≃ 2.74 Hz versus *v_D_* ≃ 20.67 Hz. Interestingly, the main macroscopic differences between the two distributions arises at short time intervals. For *τ_α_* = 2 ms, (see Fig. 4(b)) an isolated and extremely narrow peak appears at ISI_0_. This first peak corresponds to the supra-threshold tonic-firing of the isolated neuron, as reported above. After this first peak, a gap is clearly visible in the *P*(*ISI*) followed by an exponential tail. The origin of the gap resides in the fact that ISI_0_ >> *τ_α_*, because if the neuron is firing tonically with its period ISI_0_ and receives a single PSP, the membrane potential has time to decay almost to the reset value V_r_ before the next spike emission. Thus a single PSP will delay the next firing event by a fixed amount corresponding to the gap in Fig. 4(b). Indeed one can estimate analytically this delay due to the arrival of a single *α*-pulse, in the present case this gives ISI_1_ = 15.45 ms, in very good agreement with the results in Fig. 4(b). No further gaps are discernible in the distribution, because it is highly improbable that the neuron will receive two (or more) PSPs exactly at the same moment at reset, as required to observe further gaps. The reception of more PSPs during the ramp up phase will give rise to the exponential tail in the *P*(*ISI*). In this case the contribution to the *CV* comes essentially from this exponential tail, while the isolated peak at ISI_0_ has a negligible contribution.

On the other hand, if *τ_α_* > ISI_0_, as in the case reported in Fig. 4(a), *P*(*ISI*) does not show anymore a gap, but instead a continuous distribution of values. This because now the inhibitory effects of the received PSPs sum up, leading to a continuous range of delayed firing times of the neuron. The presence of this peak of finite width at short *ISI* in the *P*(*ISI*) plus the exponentially decaying tail are at the origin of the observed *CV* > 1. In Fig. 4 (e) and 4 (f) the distributions of the coefficient 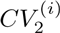 are also displayed for the considered neurons as black lines with symbols. These distributions clearly confirm that the dynamics are bursting for the longer synaptic time scale and essentially Poissonian for the shorter one.

We would like to understand whether it is possible to reproduce similar distributions of the ISIs by considering an isolated cell receiving Poissonian distributed inhibitory inputs. In order to verify this, we simulate a single cell receiving *K* uncorrelated spike trains at a rate ⟨*ν*⟩*_N_*, or equivalently, a single Poissonian spike train with rate *K*⟨*ν*⟩*_N_*. Here, ⟨*ν*⟩*_N_* is the average firing rate of a single neuron in the original network. The corresponding *P*(*ISI*) are plotted in Fig. 4 (c) and 4 (d), for *τ_α_* = 20 ms and 2 ms, respectively. There is a remarkable similarity between the reconstructed ISI distributions and the real ones (shown in Fig. 4(a) and (b)) , in particular at short ISIs. Also the distributions of the 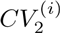 for the reconstructed dynamics are similar to the original ones, as shown in Fig. 4 (e) and 4 (f). Altogether, these results demonstrate that the bursting activity of coupled inhibitory cells is not a consequence of complex correlations among the incoming spike trains, but rather a characteristic related to intrinsic properties of the single neuron: namely, its tonic firing period, the synaptic strength, and the post-synaptic time decay. The fundamental role played by long synaptic time in inducing bursting activity has been reported also in a study of a single LIF neuron below threshold subject to Poissonian trains of exponentially decaying PSPs [32].

Obviously this analysis cannot explain collective effects, like the non trivial dependence of the number of active cells *n*\* on the synaptic strength, discussed in the previous sub-section, or the emergence of correlations and anti-correlations among neural assemblies (measured by *σ*(*C*)) due to the covarying of the firing rates in the network, as seen in striatal slices and shown in Fig. 1 (c) for our model. To better investigate the influence of *τ_α_* on the collective properties of the network we report in Fig. S4(a) and (b) the averaged *CV*, *σ*(*C*), *n*\* and Δ*M_d_* for *τ_α_* ϵ [2, 50] ms. As already noticed, the network performs better in mimicking the MSN dynamics and in discriminating between different inputs at larger *τ_α_* (e.g. at 20 ms). However, what is the minimal value of *τ_α_* for which the network still reveals cell assembly dynamics and discriminative capabilities ? From the data shown in Fig. S4(a) one can observe that *σ*(*C*) and Δ*M_d_* attain their maximal values in the range 10 ms ≤ *τ_α_* ≤ 20 ms. This indicates that clear cell assembly dynamics with associated good input pattern discrimination can be observed in this range. However, the bursting activity is not particularly pronounced at *τ_α_* = 10 ms, where ⟨*CV*⟩*_N_* ≃ 1. Therefore only the choice *τ_α_* = 20 ms fulfills all the requirements.

Interestingly, genetic mouse models of Huntington’s disease (HD) revealed that spontaneuous IPSCs in MSNs has a reduced decay time and half-amplitude duration compared to wild-types [20]. Our analysis clearly indicate that a reduction of *τ_α_* results in more stochastic single-neuron dynamics, as indicated by ⟨*CV*⟩*_N_* ≃ 1, as well as in a less pronounced structured assembly dynamics (Fig. S4(a)). This resembles what observed for the striatum dynamics of freely behaving mice with symptomatic HD [21]. In particular, the authors have shown in [21] that at the single unit level HD mice reveals a *CV* ≃ 1 in contrast to *CV* ≃ 2 for wild-type mice, furthermore the level of correlation among the neural firings was definitely reduced in HD mice.

**Figure 4.**
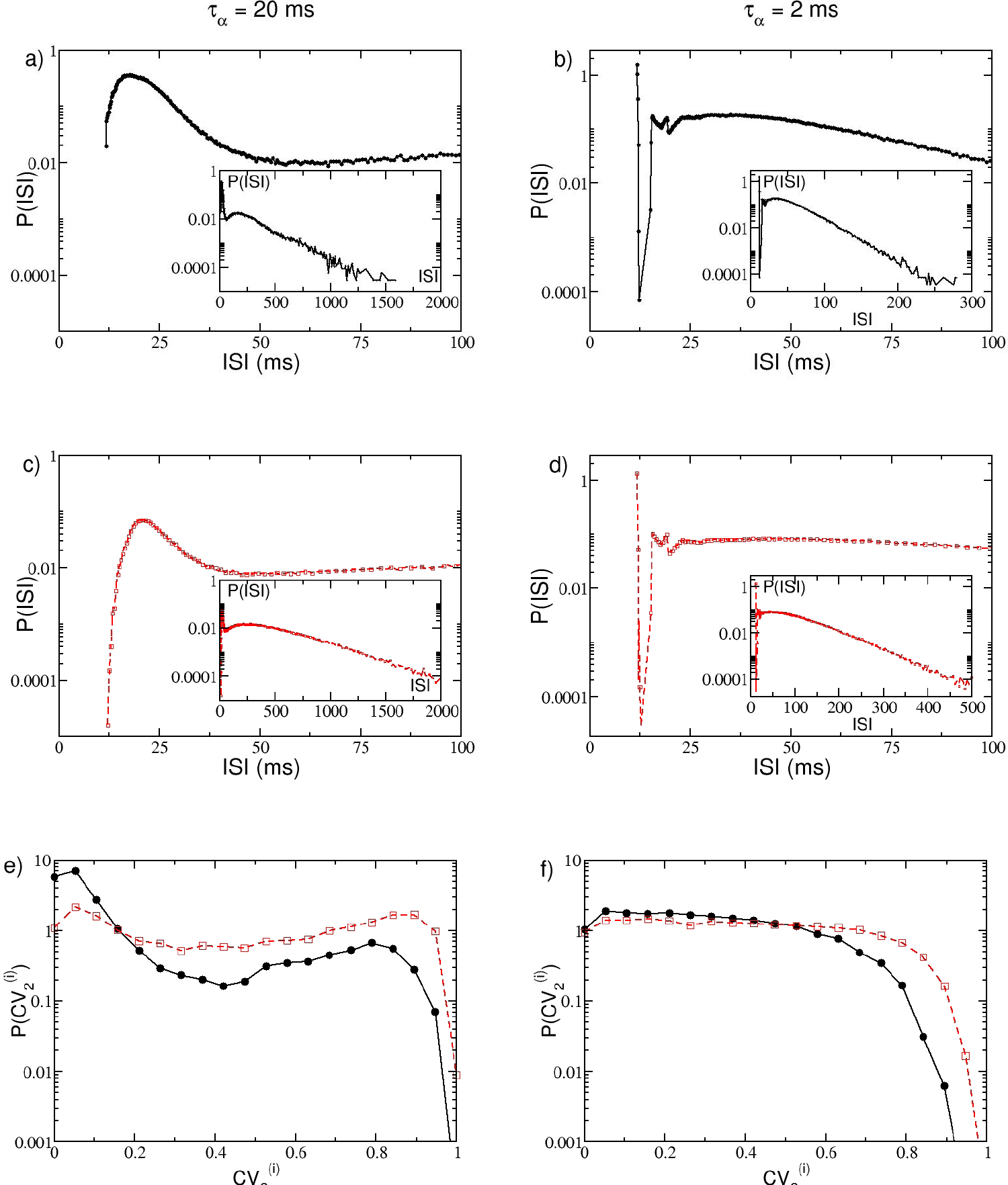
Single neuron statistics. First row: distributions *P*(*ISI*) for one representative cell in the network are shown in black. Second row: the corresponding Poissonian reconstruction of the *P*(*ISI*) are reported in red. In all plots the main figure displays the distributions at short ISIs, while the inset is a zoom out of the whole distribution. Third row: single neuron distribution of the 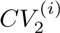 for the considered neuron (black solid lines with circles) and its Poissonian distribution (red dashed line with squares). The left (right) column corresponds to *τ_α_* = 20 (2 ms). The network parameters are Δ*V* = 5 mV and *g* = 8, and the others as in Fig. 1, both the examined neurons have *I_s_* = –45.64 mV. For the Poissonian reconstruction the frequencies of the incoming uncorrelated spike trains are set to ⟨*ν*⟩*_N_* ≈ 7.4 Hz (⟨*ν*⟩*_N_* ≈ 8.3 Hz) for *τ_α_* = 20ms (*τ_α_* = 2ms), as measured from the corresponding network dynamics. The distributions are obtained by considering a sequence of 10^9^spikes in the original network, and 10^7^ events for the Poissonian reconstruction.

### Origin of the cell assemblies

A question that we have not addressed so far is: how do cell assemblies arise ? Since the network is purely inhibitory it is reasonable to guess that correlation (anti-correlation) among groups of neurons will be related to the absence (presence) of synaptic connections between the considered groups. In order to analyze the link between the correlation and the network connectivity we compare the clustered cross-correlation matrix of the firing rates *C*(*ν_i_, ν_j_*) (shown in Fig 5 (a)) with the associated connectivity matrix *C_ij_* (reported in Fig. 5 (b)). The cross-correlation matrix is organized in *k* = 15 clusters via the *k-means* algorithm, therefore we obtain a matrix organized in a *k* χ *k* block structure, where each block (*m, l*) contains all the cross-correlation values of the elements in cluster *m* with the elements in cluster *l*. The connectivity matrix is arranged in exactly the same way, however it should be noticed that while *C*(*ν_i_, ν_j_*) is symmetric, the matrix *C_ij_* is not symmetric due to the unidirectional nature of the synaptic connections. From a visual comparison of the two figures it is clear that the most correlated blocks are along the diagonal and that the number of connections present in these diagonal blocks is definitely low, with respect to the expected value from the whole matrix. An exception is represented by the largest diagonal block which reveals, however, an almost zero level of correlation among its members. We have highlighted in blue some blocks with high level of anti-correlations among the elements, the same blocks in the connectivity matrix reveal a high number of links. A similar analysis, leading to the same conclusions was previously reported in [14].

To make this comparison more quantitative, we have estimated for each block the average cross-correlation, denoted as ⟨*C*⟩*_ml_*, and the average probability *p_ml_* of unidirectional connections from the cluster *l* to the cluster *m*. These quantities are shown in Fig. 5 (c) for all possible blocks. The correlation ⟨*C*⟩*_ml_* decreases with the probability *p_ml_*, and a linear fit to the data is reported in the figure as a solid black line. However, there are blocks that are outliers with respect to this fit. The blocks along the main diagonal (black squares) all have high correlation values ⟨*C*⟩*_mm_* and low probabilities *p_mm_*, smaller than the average probability *p* = 0.05, shown as a dashed vertical red line in Fig. 5 (c). An exception is represented by a single black square located exactly on the linear fit in proximity of *p* = 0.05 this is the large block with almost zero level of correlation among its elements previously identified. Furthermore, the blocks with higher anticorrelation, denoted as blue triangles in the figure, have probabilities *p_ml_* definitely larger than 5 %. The exceptions are two triangles lying exactly on the vertical dashed line corresponding to 5 %. This is due to the fact that the *p_ml_* are not symmetric, and it is sufficient to have a large probability of connections in only one of the two possible directions between blocks *m* and *l* to observe anti-correlated activities between the two assemblies.

Having clarified that the origin of the assemblies identified from the correlations of the firing rates is directly related to structural properties of the networks, we would like to understand if the neurons belonging to the assemblies also share other properties. In particular, we can measure the similarity of the neurons within each block (*m, l*) by estimating the block averaged similarity metrics *e_ml_* introduced in Eq. (10) in Methods. This quantity measures how similar are the effective synaptic inputs *W_i_* of two neurons in the network. In Fig. 5 (d) we report ⟨*C*⟩*_ml_* versus *e_ml_* for all the blocks. It is evident that the diagonal blocks (black squares in the figure) have a higher similarity value *e_mm_* with respect to the average (the vertical red dashed line in Fig. 5 (d)). Thus suggesting that correlated assemblies are formed by neurons with similar effective inputs and with few structural connections among them. However, the observed anticorrelations are only due to structural effects since the most anticorrelated blocks (blue triangles in Fig. 5 (d)) do not reveal any peculiar similarity value *e_ml_*. Similar results have been obtained by considering the neural excitability *I_i_* instead of the effective synaptic input *W_i_*.

**Figure 5.**
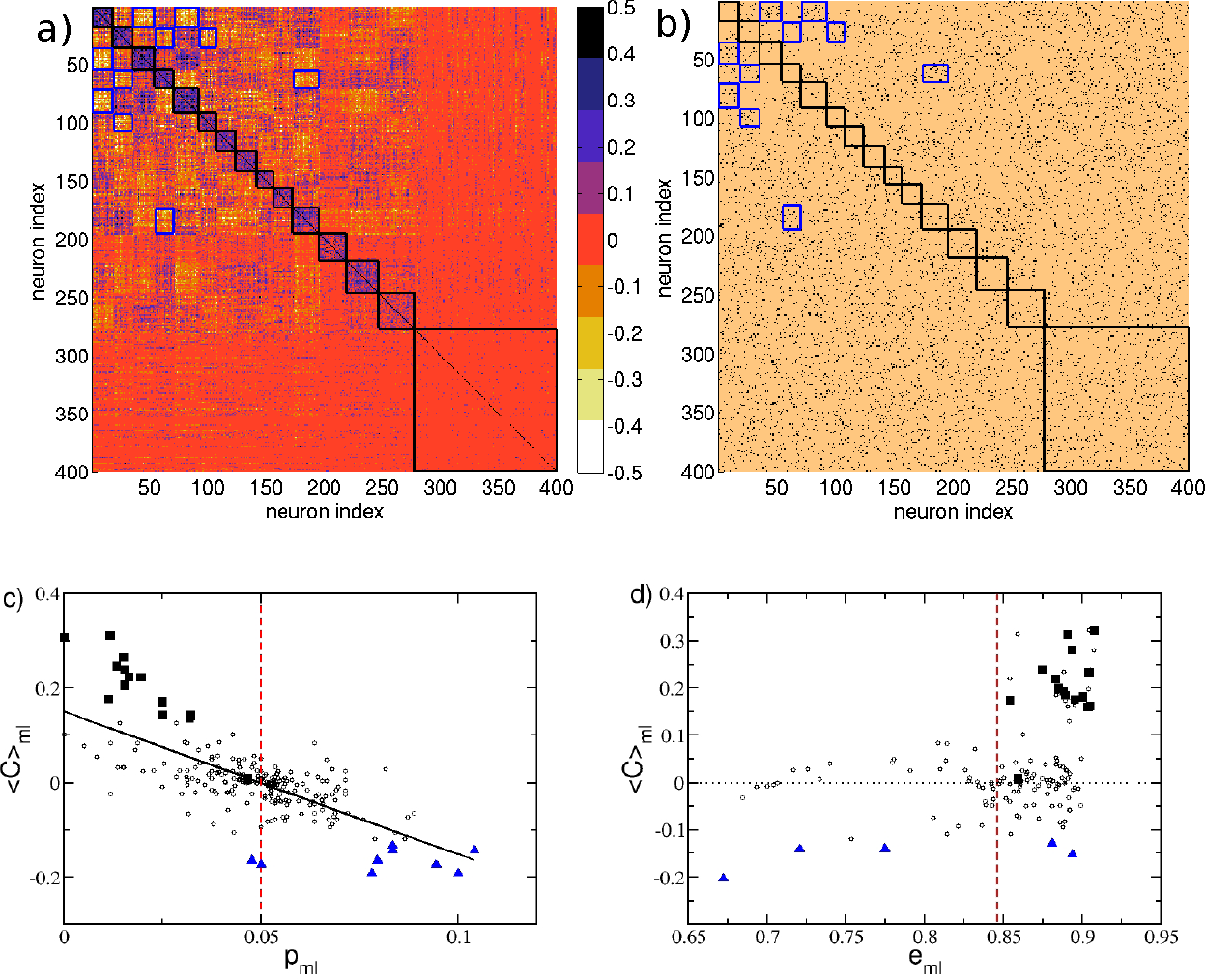
Cell assemblies and connectivity. a) Cross-correlation matrix *C*(*υ_i_, ν_j_*) of the firing rates organized according to the clusters generated via the *k-means* algorithm with *k* = 15, the clusters are ordered as in Fig. 1(c) from the highest to the lowest correlated one. b) Connectivity matrix *C_ij_* with the indices ordered as in panel a). Here, a black (copper) dot denotes a 1 (0) in *C_ij_*, i.e. the presence of a synaptic connection from *j* to *i*. c) Average cross-correlation ⟨*C*⟩*_ml_* among the elements of the matrix block (*m, l*), versus the probability *p_ml_* to have synaptic connections from neurons in the cluster l to neurons in the cluster *m*. d) ⟨*C*⟩*_ml_* versus the block averaged similarity metrics *e_ml_*. Black squares indicate the blocks along the diagonal delimited by black borders in panel a) and b); blue triangles denote the ten blocks with the lowest ⟨*C*⟩*_ml_* values, which are also delimited by blue edges in a) and b). The vertical red dashed line in panel c) denotes the average probability to have a connection *p* = 5% and in panel d) the value of the metrics *e_ml_* averaged over all the blocks. The black solid line in panel c) is the linear regression to the data (⟨*C*⟩*_ml_* ≈ 0.15 – 3.02*p_ml_*, correlation coefficient *R* = –0.72). Other parameters as in Fig. 1.

### Discriminative and computational capability

In this sub-section we examine the ability of the network to perform different tasks: namely, to respond in a reproducible manner to stimuli and to discriminate between similar inputs via distinct dynamical evolution. For this analysis we have always compared the responses of the network obtained for a set of parameters corresponding to the maximum *Q*_0_ value shown in Fig. 2(d), where *τ_α_* = 20 ms, and for the same parameters but with a shorter PSP decay time, namely *τ_α_* = 2 ms.

To check for the capability of the network to respond to cortical inputs with a reproducible sequence of states, we perform a simple experiment, following the protocol described in [15, 16], where two different inputs *I*^(1)^ and *I*^(2)^ are presented sequentially to the system. Each input persists for a time duration *T_sw_* and then the stimulus is switched to the other one and this process is repeated for the whole simulation time. The raster plot measured during such an experiment is shown in Fig. 6 (a) for *τ_α_* = 20 ms. Whenever one of the stimuli is presented, a specific sequence of activations can be observed. Furthermore, the sequence of emerging activity patterns is reproducible when the same stimulus is again presented to the system, as can be appreciated by observing the patterns encircled with black lines in Fig. 6 (a).

**Figure 6.**
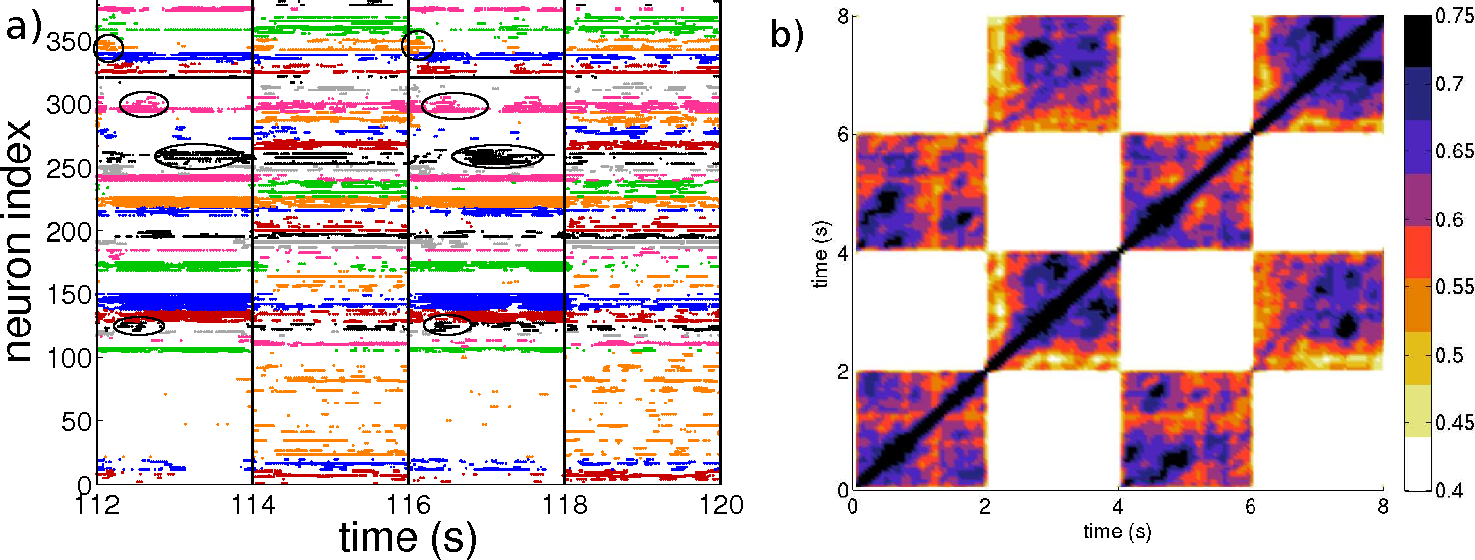
Sequential switching. a) Raster plot associated to the two input protocols *I*^(i)^ and *I*^(2)^. The circles denote the clusters of active neurons appearing repetitively after the presentation of the stimulus *I*^(i)^. Vertical lines denote the switching times between stimuli. The clustering algorithm employed to identify the different groups is applied only during the presentation of the stimulus *I*^(1)^ Itherefore the sequential dynamics is most evident for that particular stimulus. b) Averaged State Transition Matrix 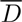, obtained by considering a 4*T_sw_* χ 4*T_sw_* sub-matrix averaged over *r* = 5 subsequent time windows of duration 4*T_sw_* (see the section *Methods* for details). The inputs *I*^(1)^ and *I*^(2)^ are different realization of the same random process, they are obtained by selecting *N* current values *I_i_* from the flat interval [*V_th_, V_th_* + Δ*V*]. The input stimuli are switched every *T_sw_* = 2 s. Number of clusters *k* = 35 in a). Other parameters as in Fig. 1.

Furthermore, we can quantitatively compare the firing activity in the network at different times by estimating the STM. Similarity is quantified by computing the normalized scalar product of the instantaneous firing rates of the *N* neurons measured at time *t_i_* and *t_j_*. We observe that the similarity of the activity at a given time *t*_0_ and at successive times *t*_0_ + 2*mT_sw_* is high (with values between 0.5 and 0.75), while it is essentially uncorrelated with the response at times corresponding to the presentation of a different stimulus, i.e. at *t*_0_ + (2*m* – 1)*T_sw_* (since the similarity is always smaller than 0.4) (here, *m* = 1, 2, 3…). This results in a STM with a periodic structure of period *T_sw_* with alternating high correlated blocks followed by low correlated blocks (see Fig. S5(b)). An averaged version of the STM calculated over a sequence of 5 presentations of *I*^(1)^ and *I*^(2)^ is shown in Fig. 6 (b) (for details of the calculation see Methods). These results show not only the capability of the network to distinguish between stimuli, but also the reproducible nature of the system response. In particular, from Fig. 6 (b) it is evident how the patterns associated with the response to the stimulus *I*^(1)^ or *I*^(2)^ are clearly different and easily identifiable. We also repeated the numerical experiment for another different realization of the inputs, noticing essentially the same features previously reported (as shown in Fig. S5(a-c)). Furthermore, to test for the presence of memory effects influencing the network response, we performed a further test where the system dynamics was completely reset after each stimulation and before the presentation of the next stimulus. This had no apparent effect in the network response.

**Figure 7.**
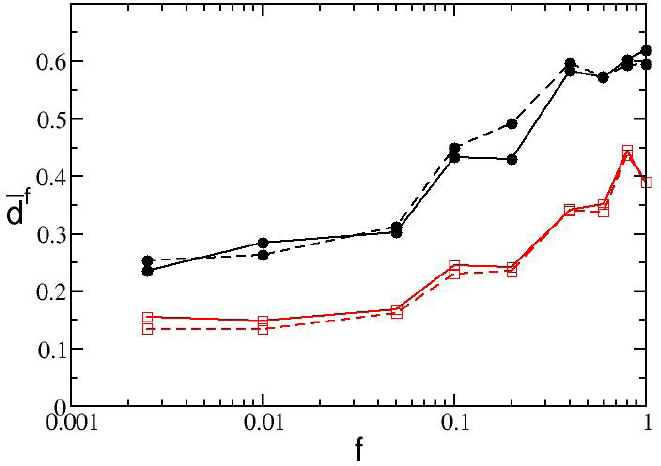
Pattern separation. Average dissimilarity as a function of the fraction *f* of inputs differing from the control input, for the values of *τ_α_* = 20ms (black circles) and *τ_α_* = 2ms (red squares) with two different observation windows *T_E_* = 2s (solid line) and *T_E_* = 10s (dashed line). Other parameters used: Δ*T* = 50ms, Δ*V* = 5 mV. Remaining parameters as in Fig. 1.

Next, we examined the influence of the PSP time scale on the observed results. In particular, we considered the case *τ_α_* = 2 ms, in this case the network (as shown in Fig. S5(d)) responds in a quite uniform manner during the presentation of each stimulus. Furthermore, the corresponding STM reported in Fig. S5(e) shows highly correlated blocks alternating with low correlated ones, but these blocks do not reveal any internal structure characteristic of cell assembly encoding.

We proceeded to check the ability of the network to discriminate among similar inputs and how this ability depends on the temporal scale of the synaptic response. In particular, we tried to answer to the following question: if we present two inputs that differ only for a fraction *f* of the stimulation currents, which is the minimal difference between the inputs that the network can discriminate ? In particular, we considered a control stimulation 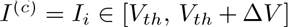 and a perturbed stimulation *I*^(*p*)^, where the stimulation currents differ only over a fraction *f* of currents *I_i_* (which are randomly chosen from the same distribution as the control stimuli). We measure the differences of the responses to the control and to the perturbed stimulations by measuring, over an observation window *T_E_*, the dissimilarity metric **d^f^** (*t*), defined in Methods. The time averaged dissimilarity metric 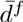 is reported as a function of *f* in Fig. 7 for two different values τ_α_. It is clear that for any *f*-value the network with longer synaptic response always discriminates better between the two different stimuli than the one with shorter PSP decay. We have also verified that the metric is robust to the modification of the observation times *T_E_*, because the dissimilarity *d^f^* (*t*) rapidly reaches a steady value (as shown in Fig. S6(a) and (b)).

To better characterize the computational capability of the network and the contribution of the PSP duration, we measured the complexity of the output signals as recently suggested in [33]. In particular, we have examined the response of the network to a sequence of three stimuli, each being a constant vector of randomly chosen currents. The three different stimuli are consecutively presented to the network for a time period *T_sw_*, and the stimulation sequence is repeated for the whole experiment duration *T_E_*. The output of the network can be represented by the instantaneous firing rates of the *N* neurons measured over a time window Δ*T* = 100 ms, this is a high dimensional signal, where each dimension is represented by the activity of a single neuron. The complexity of the output signals can be estimated by measuring how many dimensions are explored in the phase space. With greater stationarity of firing rates, fewer variables are required to reconstruct the whole output signal [33].

**Figure 8.**
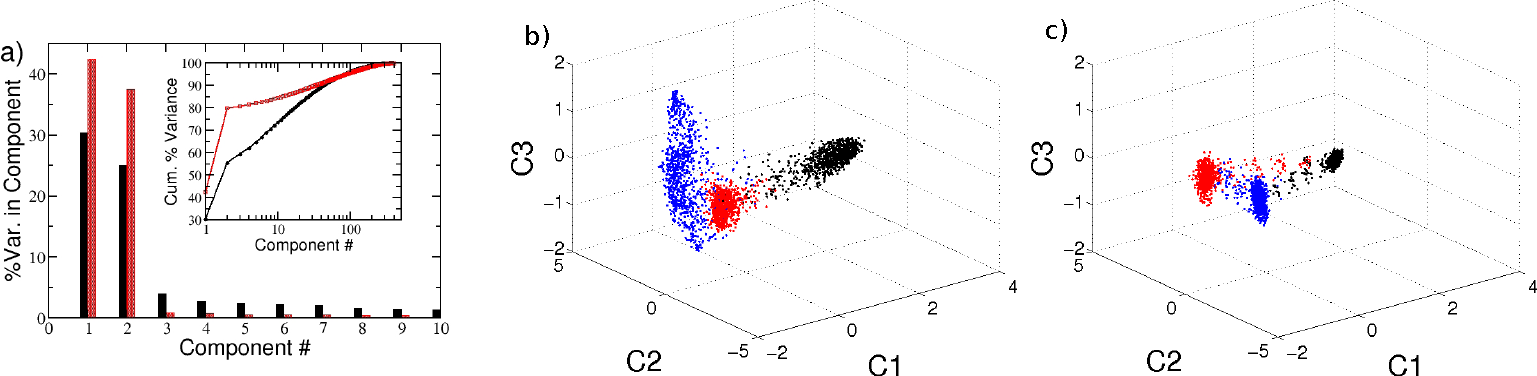
Computational capability of the network. Characterization of the firing activity of the network, obtained as response to three consecutive inputs presented in succession. a) Percentage of the variance of the neuronal firing activity reproduced by each of the first 10 principal components. The inset displays the corresponding cumulative percentage as a function of the considered component. Filled black and shaded red (bar or symbols) correspond to *τ_α_* = 20 ms and *τ_α_* = 2 ms, respectively. Projection of the neuronal response along the first three principal components for b) *τ_α_* = 20 ms and c) *τ_α_* = 2 ms. Each point in the graph correspond to a different time of observation. The three colors denote the response to the three different inputs, which are quenched stimulation currents randomly taken as 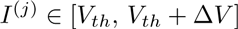 for *j* = 1, 2, 3, the experiment is then performed as explained in the text.

A principal component analysis (PCA) performed over *T_E_*/Δ*T* observations of the *N* firing rates reveals that for *τ_α_* = 2 ms 80% of the variance is recovered already with a projection over a two dimensional sub-space (red bars in Fig. 8 (a)). On the other hand, for *τ_α_* = 20 ms a higher number of principal components is required to reconstruct the dynamical evolution (black bars in Fig. 8 (a)), thus suggesting higher computational capability of the system with longer PSPs [33].

These results are confirmed by analyzing the projections of the firing rates in the subspace spanned by the first three principal components (*C*1, *C*2, *C*3) shown in Fig. 8 (b) and (c) for *τ_α_* = 20 ms and *τ_α_* = 2 ms, respectively. The responses to the three different stimuli can be effectively discriminated by both networks, since they lie in different parts of the phase space. However, the response to the three stimuli correspond essentially to three fixed points for *τ_α_* = 2 ms, while trajectories evolving in a higher dimension are associated to each constant stimulus for *τ_α_* = 20 ms.

These analyses confirm that the network parameters selected by employing the maximal *Q*_0_ criterion also result in a reproducible response to different stimuli, as well as in an effective discrimination between different inputs.

In recent work Ponzi and Wickens [16] noticed that in their model the striatally relevant regimes correspond to marginally stable dynamical evolution. In the Supporting Information Text S1 we devote the sub-section *Linear stability analysis* to the investigation of this specific point. Our conclusion is that for our model the striatally relevant regimes are definitely chaotic, but located in proximity of a transition to linearly stable dynamic (see also Fig. S3). However for inhibitory networks it is known that even linearly stable networks can display erratic dynamics (resembling chaos) due to finite amplitude perturbations [34, 35, 36, 37]. This suggests that the usual linear stability analysis, corresponding to the estimation of the maximal Lyapunov exponent [38], is unable to distinguish between regular and irregular evolution, at least for the studied inhibitory networks [37].

### Physiological relevance for biological networks under different experimental conditions

Carrillo *et al.* [11] considered a striatal network *in vitro*, which displays sporadic and asynchronous activity under control conditions. To induce spatio-temporal patterned activity they perfused the slice preparation with N-methyl-D-aspartate (NMDA) providing tonic excitatory drive and generating bursting activity [39, 40]. The crucial role of the synaptic inhibition in shaping the patterned activity in striatal dynamics was demonstrated by applying the GABA*_a_* receptor antagonist bicuculline to effectively decrease the inhibitory synaptic effect [11].

In our simple model, ionic channels and NMDA-receptors are not modeled; nevertheless it is possible to partly simulate the effect of NMDA administration by increasing the excitability of the cells in the network, and the effect of bicuculline by an effective decrease in the synaptic strength. We examined whether these assumptions lead to results similar to those reported in [11].

In our model the single cell excitability is controlled by the parameter *I_i_*. The computational experiment consists in setting the system in a low firing regime corresponding to the control conditions with 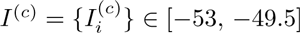 mV and in enhancing, after 20 seconds, the system excitability to the range of values 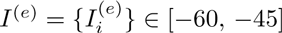 mv, for another 20 seconds. This latter stage of the numerical experiment corresponds to NMDA perfusion in the brain slice experiment. The process is repeated several times and the resulting raster plot is coarse grained as explained in Methods (sub-section *Synchronized Event Transition Matrix*).

From the coarse grained version of the raster plot, we calculate the *Network Bursting Rate* (NBR) as the fraction of neurons participating in a burst event in a certain time window. Whenever the instantaneous NBR is larger than the average NBR plus two standard deviations, this is identified as a *synchronized bursting event* (as shown in Fig. 9(a) and (f)). In Fig. 9(b) we plot all the neurons participating in a series of *S_s_* = 20 synchronized bursting events. Here the switching times between control conditions and the regimes of increased excitability are marked by vertical dashed lines. Due to the choice of the parameters, the synchronized events occur only during the time intervals during which the network is in the enhanced excitability regime. Each synchronized event is encoded in a binary *N* dimensional vector *W_s_* (*t*) with 1 (0) entries indicating that the corresponding neuron was active (inactive) during such event. We then measure the similarity among all the events in terms of the *Synchronized Event Transition Matrix* (SETM) shown in Fig. 9(c). The SETM is the normalized scalar product of all pairs of vectors *W_s_* registered in a given time interval (for more details see Methods). Furthermore, using the SETM we divide the synchronized events into clusters according to an optimal clustering algorithm [41] (see Methods). In the present case we identified 3 distinct *states* (clusters): if we project the vectors *W_s_*, characterizing each single synchronized event on the two dimensional space spanned by the first two principal components (*C*1, *C*2), we observe a clear division among the 3 states (see Fig. 9(d)). It is now important to understand whether the cells firing during the events classified as a state are the same or not. We observe that the groups of neurons recruited for each synchronized event corresponding to a certain state largely overlap, while the number of neurons participating to different states is limited. As shown in Fig. 9(e), the number of neurons participating in a certain state is of the order of 40-50, while the coactive neurons (those participating in more than one state) ranges from 12 to 25. Furthermore, we have a core of 9 neurons which are firing in all states. Thus we can safely identify a distinct assembly of neurons active for each state.

As shown in Fig. 9(c), we observe that the system alternates its activity among the previously identified cell assemblies. In particular, we have estimated the transition probabilities from one state to any of the three identified states. We observe that the probability to remain in state 2 or to arrive to this state from state 1 or 3 is quite high, ranging between 38 and 50 %, therefore this is the most visited state. The probability that two successive events are states of type 1 or 2 is also reasonably high ranging from ≃ 29 – 38% as well as the probability that from state 1 one goes to 2 or viceversa (≃ 38 – 43%). Therefore the synchronized events are mostly of type 1 and 2, while the state 3 is the less *attractive*, since the probability of arrving to this state from the other ones or to remain on it once reached, are between 25 - 29 %. If we repeat the same experiment after a long simulation interval *t* ≃ 200 s we find that the dynamics can be always described in terms of small number of states (3-4), however the cells contributing to these states are different from the ones previously identified. This is due to the fact that the dynamics is in our case chaotic, as we have verified in the Supporting Information Text S1 (*Linear Stability Analysis*). Therefore even small differences in the initial state of the network, can have macroscopic effects on sufficiently long time scales.

To check for the effect of bicuculline, the same experiment is performed again with a much smaller synaptic coupling, namely *g* = 1, the results are shown in Fig. 9(f-j). The first important difference can be identified in higher NBR values with respect to the previous analyzed case (*g* = 8) Fig. 9(f). This is due to the decreased inhibitory effect, which allows most of the neurons to fire almost tonically, and therefore a higher number of neurons participate in the bursting events. In Fig. 9(g) a highly repetitive pattern of synchronized activity (identified as state 2, blue symbols) emerges immediately after the excitability is enhanced. After this event we observe a series of bursting events, involving a large number of neurons (namely, 149), which have been identified as an unique cluster (state 1, red symbols). The system, analogously to what shown in [11], is now locked in an unique state which is recurrently visited until the return to control conditions. Interestingly, synchronized events corresponding to state 1 and state 2 are highly correlated when compared with the *g* = 8 case, as seen by the SETM in Fig. 9(h). Despite this, it is still possible to identify both states when projected on the two dimensional space spanned by the first two principal components (see Fig. 9(i)). This high correlation can be easily explained by the fact that the neurons participating in state 2 are a subset of the neurons participating in state 1, as shown in Fig. 9(j). Furthermore, the analysis of the transition probabilities between states 1 and 2 reveals that starting from state 2 the system never remains in state 2, but always jumps to state 1. The probability of remaining in state 1 is high ≃ 64%. Thus we can affirm that in this case the dynamics are dominated by the state 1.

To determine the statistical relevance of the results presented so far, we repeated the same experiment for ten different random realizations of the network. The detailed analysis of two of these realizations is reported in Figs. S7(a-h) (see also Text S1). We found that, while the number of identified states may vary from one realization to another, the consistent characteristics that distinguish the NMDA perfused scenario and the decreased inhibition one, are the variability in the SETM and the fraction of coactive cells. More precisely, on one hand the average value of the elements of the SETM is smaller for *g* = 8 with respect to the *g* = 1 case, namely 0.54 versus 0.84, on the other hand their standard deviation is larger, namely 0.15 versus 0.07. This indicates that the states observed with *g* = 1 are much more correlated with respect to the states observable for *g* = 8, which show a larger variability. The analysis of the neurons participating to the different states revealed that the percentage of neurons coactive in the different states passes from 51 % at *g* = 8 to 91 % at *g* = 1. Once more the reduction of inhibition leads to the emergence of states which are composed by almost the same group of active neurons, representing a dominant state. These results confirm that inhibition is fundamental to cell assembly dynamics.

Altered intrastriatal signaling has been implicated in several human disorders, and in particular there is evidence for reduced GABAergic transmission following dopamine depletion [42], as occurs in Parkinson’s disease. Our simulations thus provide a possible explanation for observations of excessive entrainment into a dominant network state in this disorder [43, 23].

## Discussion

In summary, we have shown that lateral inhibition is fundamental for shifting the network dynamics from a situation where a few neurons, tonically firing at a high rate, depress a large part of the network, to a situation where all neurons are active and fire with similar slow rates. In particular, if inhibition is too low, or too transient, a *winner takes all* mechanism is at work and the activity of the network is mainly mean-driven. By contrast, if inhibition has realistic strength and duration, almost all the neurons are on average sub-threshold and the dynamical activity is fluctuation-driven [30].

Therefore we can reaffirm that the MSN network is likely capable of producing slow, selective, and reproducible activity sequences as a result of lateral inhibition. The mechanism at work is akin to the *winerless competition* reported to explain the function of olfactory networks for the discrimination of different odors [44]. Winnerless competition refers to a dynamical mechanism, initially revealed in asymmetrically coupled inhibitory rate models [45], displaying a transient slow switching evolution along a series of metastable saddles (for a recent review on the subject see [46]). In our case, the sequence of metastable states can be represented by the firing activity of the cell assembly, switching over time. In particular, slow synapses have been recognized as a fundamental ingredient, along with asymmetric inhibitory connections, for observing the emergence of winnerless competition in realistic neuronal models [47, 48].

We have introduced a new metric to encompass in a single indicator key aspects of this patterned sequential firing, and with the help of this metric we have identified the parameter ranges for best obtaining these dynamics. Furthermore, for these parameters the network is able to respond in a reproducible manner to the same stimulus, presented at different times, while presenting complex computational capability by responding to constant stimuli with an evolution in a high dimensional space [33].

Our analysis confirms that the IPSP/IPSC duration is crucial in order to observe bursting dynamics at the single cell level as well as structured assembly dynamics at the population level. A reduction of the synaptic time has been observed in symptomatic HD mice [20]. In our model this reduction leads single neurons towards a Poissonian behaviour and to a reduced level of correlation/anticorrelation among neural assemblies, in agreement with experimental results reported for mouse models of HD [21].

In summary, we have been able to reproduce general experimental features of MSN networks in brain slices [11]. In particular, we observed a structured activity alternating among a small number of distinct cell assemblies. Furthermore, we have reproduced the dynamical effects induced by decreasing the inhibitory coupling: the drastic reduction of the inhibition leads to the emergence of a dominant highly correlated neuronal assembly. This may help account for the dynamics of Parkinsonian striatal microcircuits, where dopamine deprivation impairs the inhibitory feedback transmission among MSNs [42, 23]. Network models such as the one presented here offer a path towards understanding just how pathologies that affect single neurons lead to aberrant network activity patterns, as seen in Parkinson’s and Huntington’s diseases, and this is an exciting direction for future research.

**Figure 9.**
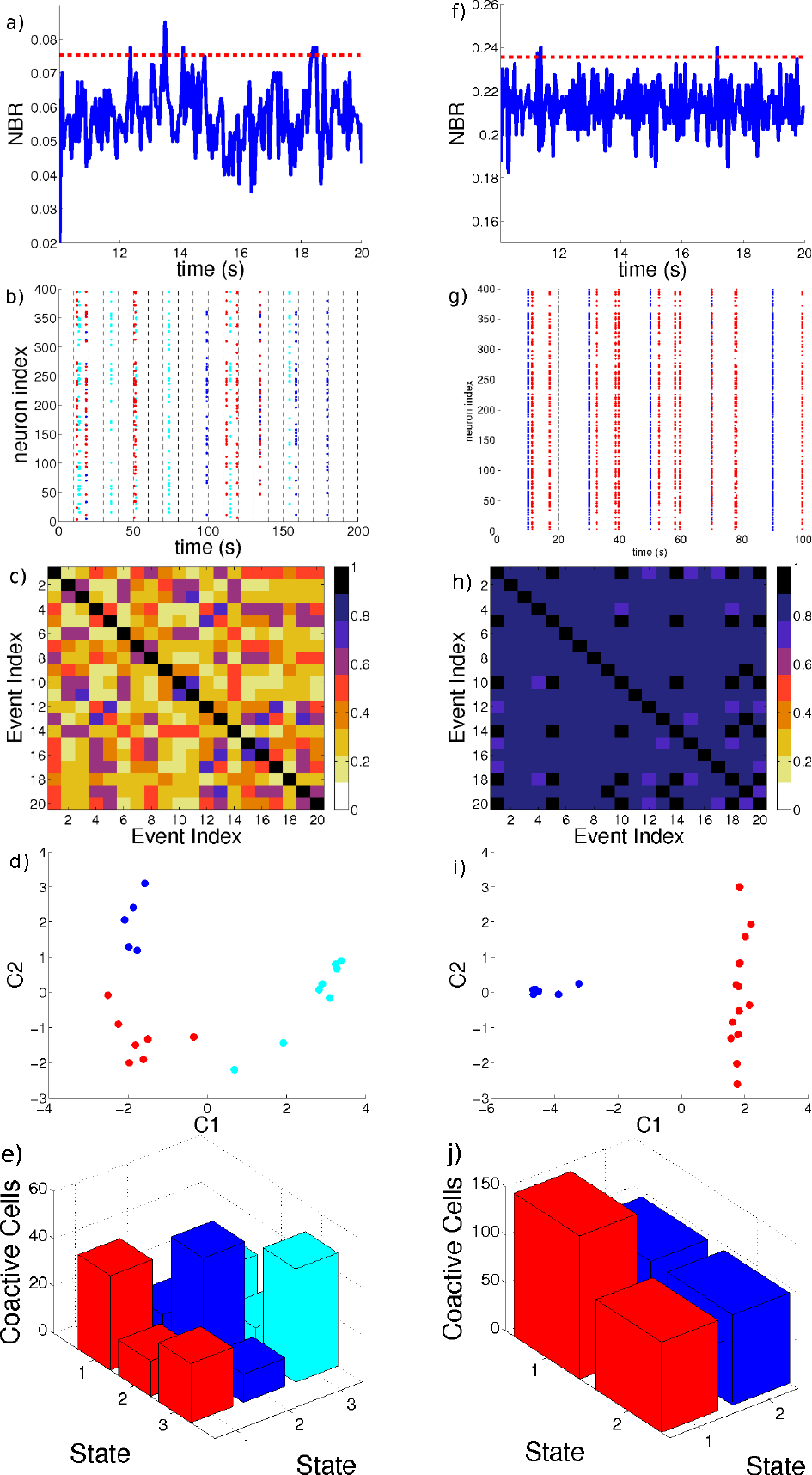
Response of the network to an increase in the excitability. a,f) Network Bursting Rate, and the threshold defined for considering a synchronized event. b,g) Neurons involved in the synchronized events, vertical lines denoted the switching times between the excited *I*^(*e*)^ and control *I*^(*c*)^ inputs. Colors in the raster indicates the group assigned to the synchronous event using an optimal community partition algorithm. c,h) Synchronized Event Transition Matrix, calculated with a window *T_W_* = 50 ms and number of events *S_s_* = 20. d,i) Projection of the synchronized events in the 2D space spanned by the first two principal components associated to the covariance matrix of the vectors *W_s_*. e,j) Number of coactive cells in each state. The diagonal elements of the bar graph represents the total number of neurons contributing to one state. Panels (a-e) correspond to *g* = 8, while panels (f-j) depict the case *g* = 1. In both cases the system is recorded during the time span required to identify *S_s_* = 20. Remaining parameters as in Fig. 1.

## Methods

### The model

We considered a network of N LIF inhibitory neurons coupled via α pulses, which can be represented via the following set of 3*N* equations.

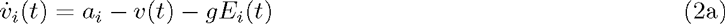

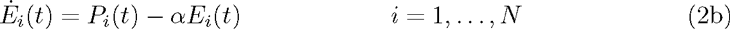

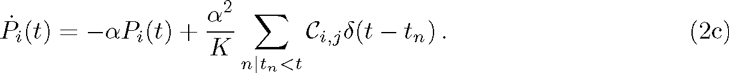

In this model *v_i_* is the membrane potential of neuron *i, K* denotes the number of pre-synaptic connections, *g* > 0 is the strength of the synapses, the variable *E_i_* accounts for the past history of previous recurrent post-synaptic potentials (PSP) that arrived to the neuron *i* at times *t_n_*, and *P_i_* is an auxiliary variable. *a_i_* is the external excitation and α is inversely proportional to the decaying time of the PSP. The inhibition is introduced via the minus sign in front of the synaptic strength in Eq. (2a). The matrix *C_i,j_* is the connectivity matrix where entry *i, j* is equal to 1 if there exists a synaptic connection from neuron *j* to neuron *i*. When the membrane potential of the *q*-th neuron arrives to the threshold *v_th_* = 1, it is reset to the value *v_r_* = 0 and the cell emits an α-pulse *p_α_*(*t*) = α^2^*t* exp (–*αt*) which is instantaneously transmitted to all its post-synaptic neurons. The α-pulses are normalized to one, therefore the area of the transmitted PSPs is conserved by varying the parameter α.

The equations (2a) to (2c) can be exactly integrated from the time *t = t*_*n*_, just after the delivery of the n-th pulse, to time *t = t*_*n* + 1_ corresponding to the emission of the (*n* + 1)-th spike, thus obtaining an *event driven map* [27, 49] which reads as

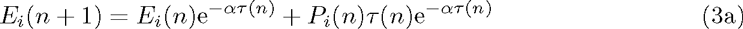

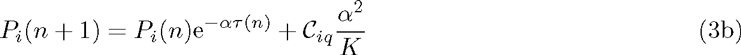

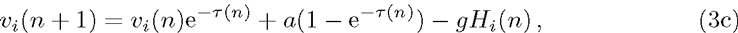

where *τ*(*n*) = *t*_*n* + 1_ – *t_n_* is the inter-spike interval associated with two successive neuronal firing in the network, which can be determined by solving the transcendental equation

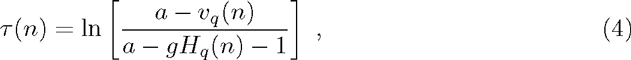

here *q* identifies the neuron which will fire at time *t*_*n* + 1_ by reaching the threshold value *v_q_*(*n* +1) = 1.

The explicit expression for *H_i_*(*n*) appearing in equations (3c) and (4) is

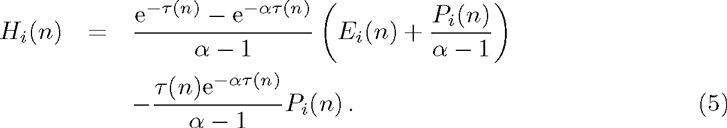

The model is now rewritten as a discrete-time map with 3*N* – 1 degrees of freedom, since one degree of freedom 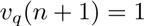, is lost due to the event driven procedure, which corresponds to performing a Poincaré section at any time a neuron fires [49].

The model so far introduced contains only adimensional units, however, the evolution equation for the membrane potential (2) can be easily re-expressed in terms of dimensional variables as follows

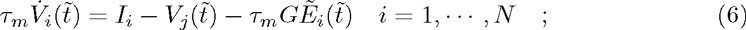

where we have chosen *τ_m_* = 10 ms as the membrane time constant in agreement with the values reported in literature for MSNs in the up state in rodents [50, 51, 52], *I_i_* represents the neural excitability and the external stimulations, which takes in account the cortical and thalamic inputs received by the striatal network. Furthermore, 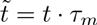, the field 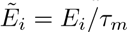 has the dimensionality of frequency and *G* of voltage.

The currents {*I_i_*} have also the dimensionality of a voltage, since they include the membrane resistance.

For the other parameters/variables the transformation to physical units is simply given by

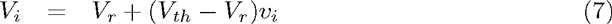

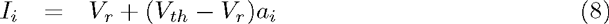

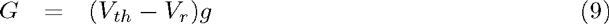

where *V_r_* = –60 mV, *V_th_* = –50 mV. The isolated *i*-th LIF neuron is supra-threshold whenever *I_i_* > *V_th_*, however due to the inhibitory coupling the effective input is 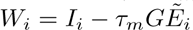. Therefore, the neuron is able to deliver a spike if 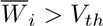, in this case the firing of the neuron can be considered as mean-driven. However, even if 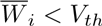, the neuron can be lead to fire from fluctuations in the effective input and the firing is in this case fluctuation-driven. It is clear that the fluctuations *σ*(*W_i_*) are directly proportional to the strength of the inhibitory coupling for constant external currents *I_i_*. The dynamics of two neurons will be equivalent whenever they have equal time averaged effective inputs 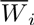. In order to measure the similarity of their dynamics we introduce the following metrics

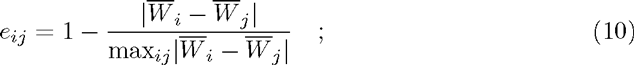

this quantity will be bounded between one and zero: the one (zero) denoting maximal (minimal) similarity.

For the PSPs the associated time constant is 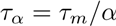, and the peak amplitude is given by

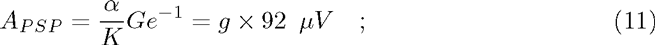

where the last equality allows for a direct transformation from adimensional units to dimensional ones, for the connectivity considered in this paper, namely *K* = 20, and for α = 0. 5, which is the value mainly employed in our analysis. The experimentally measured peak amplitudes of the inhibitory PSPs for spiny projection neurons ranges from ≃ 0.16 – 0.32 mV [7] to ≃ 1 – 2 mV [28]. These values depend strongly on the measurement conditions a renormalization of all the reported measurements nearby the firing threshold gives for the PSP peak ≃ 0.17 – 0.34 mV [9]. Therefore from Eq. (11) one can see that realistic values for *A_PSP_* can be obtained in the range *g* ∈ [2 : 10]. For α = 0.5 one gets *τ_α_* = 20 ms, which is consistent with the PSP duration and decay values reported in the literature for inhibitory transmission among spiny neurons [7, 31]

Our model does not take in account the depolarizing effects of inhibitory PSPs for *V* ≤ *E_cl_* [28]. The GABA neurotransmitter has a depolarizing effect in mature projection spiny neurons, however this depolarization does not lead to a direct excitation of the spiny neurons. Therefore our model can be considered as encompassing only the polarizing effects of the PSPs for *V* > *E_cl_*. This is the reason we have assumed that the membrane potential varies in the range [–60 : –50] mV, since *E_cl_* ≃ –60 mV and the threshold is ≃ –50 mV [28].

In the paper we have always employed dimensional variables (for simplicity we neglect the tilde on the time variable), apart from the amplitude of the synaptic coupling, for which we have found more convenient to use the adimensional quantity *g*.

### Characterization of the network activity

We define active neurons, as opposed to silent neurons, as cells that deliver a number of spikes larger than a certain threshold *S*_Θ_ =3 during the considered numerical experiments. In particular, we show in Fig. S1 of the Supporting Information that the value of the chosen threshold does not affect the reported results for 0 ≤ *S*_Θ_ ≤ 100.

Characterization of the dynamics of the active neurons is performed via the coefficient of variation *CV*, the local coefficient of variation *CV*_2_ and the zero lag cross-correlation matrix of the firing rates *C*(*υ_i_, υ_j_*). The coefficient of variation associated to the i-th neuron is then defined as the ratio:

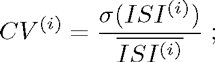

where σ(*A*) and 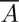 denotes the standard deviation and mean value of the quantity *A*. The distribution of the coefficient of variation *P*(*CV*) reported in the article refers to the values of the *CV* associated with all the active cells in the network.

Another useful measure of spike statistics is the local coefficient of variation. For each neuron *i* and each spike emitted at time 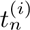 from the considered cell the local coefficient of variation is measured as

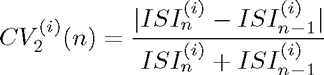

where the *n*-th inter-spike interval is defined as 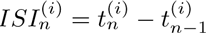. The above quantity ranges between zero and one: a zero value corresponds to a perfectly periodic firing, while the value one is attained in the limit 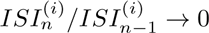 (or 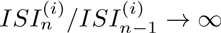). The probability distribution function *P*(*CV*_2_) is then computed by employing the values of the 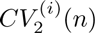 for all the active cells of the network estimated at each firing event.

The level of correlated activity between firing rates is measured via the cross-correlation matrix *C*(*υ_i_, υ_j_*). The firing rate *ν_i_*(*t*) of each neuron *i* is calculated at regular intervals Δ*T* = 50 ms by counting the number of spikes emitted in a time window of 10Δ*T* = 500 ms, starting from the considered time *t*. For each pair of neuron *i* and *j* the corresponding element of the *N* χ *N* symmetric cross-correlation matrix *C*(*i, j*) is simply the Pearson correlation coefficient measured as follows

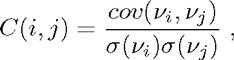

where *cov*(*v_i_, v_j_*) is the covariance between signals *v_i_*(*t*) and *v_j_*(*t*). For statistical consistency this is always calculated for spike trains containing 10^7^ events. This corresponds to time intervals *T_E_* ranging from 50 s to 350 s (from 90 s to 390 s) for Δ*V* = 5 mV (Δ*V* = 1 mV) and *g* ∈ [0.1,12]. We have also verified that the indicators entering in the definition of the metrics *Q*_0_, namely *n*\*, the average coefficient of variation ⟨*CV*⟩*_N_* and *σ*(*C*), do not depend on the duration of the considered time windows, provided these are sufficiently long. Namely, we observe that asymptotic values are already obtained for spike trains containing more than 50,000 events. This amounts to *T_E_* ranging between 250 ms and 2 s for the considered parameter values.

### State Transition Matrix (STM) and measure of dissimilarity

The STM is constructed by calculating the firing rates *v_i_*(*t*) of the *N* neurons at regular time intervals Δ*T* = 50 ms. At each time *t* the rates are measured by counting the number of spikes emitted in a window 2Δ*T*, starting at the considered time. Note that the time resolution here used is higher than the one employed for the cross-correlation matrix, since we are interested in the response of the network to a stimulus presentation evaluated on a limited time window. The firing rates can be represented as a state vector 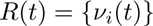 with *i* = 1, …, *N*. For an experiment of duration *T_E_*, we have 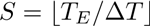 state vectors *R*(*t*) representing the network evolution (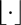 denotes the integer part). In order to measure the similarity of two states at different times *t_m_* = *m* χ Δ*T* and *t_n_* = *n* χ Δ*T*, we have calculated the normalized scalar product

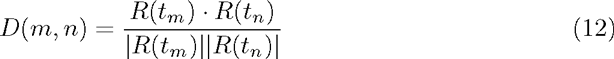

for all possible pairs *m, n* = 1, …, *S* during the time experiment *T_E_*. This gives rise to a *S χ S* matrix called the state transition matrix [53].

In the case of the numerical experiment with two inputs reported in the section *Results*, the obtained STM has a periodic structure of period *T_sw_* with high correlated blocks followed by low correlated ones (see Figs. S5(b) and (e) for the complete STM). In Fig 6 (b) we report a coarse grained version of the entire STM obtained by taking a 4*T_sw_* χ 4*T_sw_* block from the STM, where the time origin corresponds to the onset of one of the two stimuli. The block is then averaged over *r* subsequent windows of duration 4*T_w_*, whose origin is shifted each time by 2*T_sw_*. More precisely the averaged 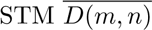 is obtained as follows:

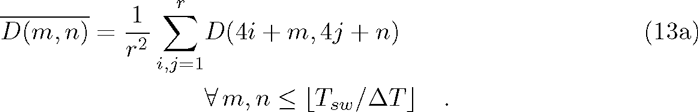

In a similar manner, we can define a dissimilarity metric to distinguish between the response of the network to two different inputs. We define a control input 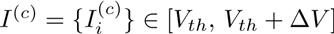, and we register the network state vectors *R^c^*(*t*) at *S* regular time intervals for a time span *T_E_*. We repeat the numerical experiment by considering the same network realization and the same initial conditions, but with a new input 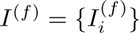. The external inputs 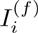 coincide with the control ones, apart from a fraction *f* which is randomly taken from the interval 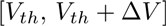. For the modified input we register another sequence *R^f^*(*t*) of state vectors on the same time interval, with the same resolution. The instantaneous dissimilarity *d^f^*(*t*) between the response to the control and perturbed stimuli is defined as:

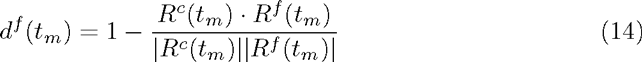

its average in time is simply given by 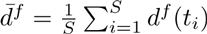. We have verified that the average 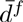 is essentially not modified if the instantaneous dissimilarities *d^f^*(*t*) are evaluated by considering the state vectors *R^c^*(*t_i_*) and *R^f^*(*t_k_*) taken at different times within the interval [0, *t_s_*] and not at the same time as done in (14).

### Distinguishability metric *Q_d_*

Following [16] a metric of the ability of the network to distinguish between two different inputs Δ*M_d_* can be defined in terms of the STM. In particular, let us consider the STM obtained for two inputs *I*^(1)^ to *I*^(2)^, each presented for a time duration *T_sw_*. In order to define Δ*M_d_* the authors in [16] considered the correlations of the state vector *R* taken at a generic time *t_m0_* with all the other configurations, with reference to Eq. (12) this amounts to examine the elements *D*(*m_0_, n*) of the STM 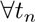. By defining *M*_1_ (*M*_2_) as the average of *D*(*m_0_, n*) over all the times *t_n_* associated to the presentation of the stimulus *I*^(1)^ (*I*^(2)^), a distinguishablity metric between the two inputs can be finally defined as

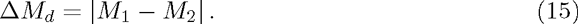

In order to take in account the single neuron variability and the number of active neurons involved in the network dynamics we have modified Δ*M_d_* by multiplying this quantity by the fraction of active neurons and the average coefficient of variation, as follows

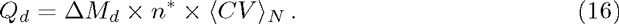

### Synchronized Event Transition Matrix (SETM)

In order to obtain a Synchronized Event Transition Matrix (SETM), we first coarse grain the raster plot of the network. This is done by considering a series of windows of duration *T_W_* = 50 ms covering the entire raster plot. A bursting event is identified whenever a neuron *i* fires 3 or more spikes within the considered window. To signal the burst occurrence a dot is drawn at the beginning of the window. From this coarse grained raster plot we obtain the Network Bursting Rate (NBR) by measuring the fraction of neurons that are bursting within the considered window. When this fraction is larger or equal to the average NBR plus two standard deviations, a synchronized event is identified. Each synchronized event is encoded in the synchronous event vector *W_s_*(*t*), a *N* dimensional binary vector where the *i*-th entry is 1 if the *i*-th neuron participated in the synchronized event and zero otherwise. To measure the similarity between two synchronous events, we make use of the normalized scalar product between all the pairs of vectors *W_s_* obtained at the different times *t_i_* and *t_j_* in which a synchronized event occurred. This represents the element *i, j* of the SETM.

### Principal Components Analysis (PCA)

In the sub-section *Discriminative and computational capability*, a Principal Component Analysis (PCA) is performed by collecting *S* state vectors *R*(*t*), measured at regular intervals Δ*T* for a time interval *T_E_*, then by estimating the covariance matrix *cov*(*v_i_, v_j_*) associated to these state vectors. Similarly, in the sub-section *Physiological relevance for biological networks under different experimental conditions* the PCA is computed by collecting the *S_s_* synchronous event vectors *W_s_*, and the covariance matrix calculated from this set of vectors.

The principal components are the eigenvectors of theses matrices, ordered from the largest to the smallest eigenvalue. Each eigenvalue represents the variance of the original data along the corresponding eigendirection. A reconstruction of the original data set can be obtained by projecting the state vectors along a limited number of principal eigenvectors.

### Clustering algorithms

The *k-means* algorithm is a commonly-used clustering technique in which *N* data points of dimension *M* are organized in clusters as follows. As a first step a number *k* of clusters is defined a-priori, then from a sub-set of the data *k* samples are chosen randomly. From each sub-set a centroid is defined in the *M*-dimensional space. at a second step, the remaining data are assigned to the closest centroid according to a distance measure. After the process is completed, a new set of *k* centroids can be defined by employing the data assigned to each cluster. The procedure is repeated until the position of the centroids converge to their asymptotic value.

An unbiased way to define a partition of the data can be obtained by finding the optimal cluster division [41]. The optimal number of clusters can be found by maximizing the following cost function, termed *modularity*:

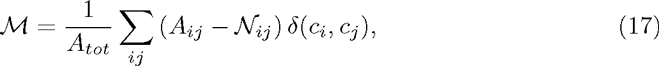

where, 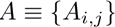 is the matrix to be clusterized, the normalization factor is 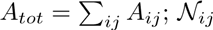 accounts for the matrix element associated to the *null model; c_i_* denotes the cluster to which the *i*-th element of the matrix belongs to, and δ(*i, j*) is the Kronecker delta. In other terms, the sum appearing in Eq. (17) is restricted to elements belonging to the same cluster. In our specific study, A is the *similarity matrix* corresponding to the SETM previously introduced. Furthermore, the elements of the matrix *N* are given by 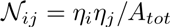, where 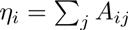, these correspond to the expected value of the similarity for two randomly chosen elements [54, 55]. If two elements are similar than expected by chance, this implies that *A_ij_ > N_ij_*, and more similar they are larger is their contribution to the modularity *M*. Hence they are likely to belong to the same cluster. The problem of modularity optimization is NP-hard [56], nevertheless some heuristic algorithms have been developed for finding local solutions based on greedy algorithms [57, 58, 59, 60]. In particular, we make use of the algorithm introduced for connectivity matrices in [61, 54], which can be straightforwardly extended to similarity matrices by considering the similarity between two elements, as the weight of the link between them [62]. The optimal partition technique is used in the sub-section *Physiological relevance for biological networks under different experimental conditions*, where it is applied to the similarity matrix 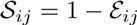 where the distance matrix 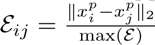. Here 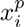 is the vector defining the *i^th^* synchronized event projected in the first *p* principal components, which accounts for the 80% of the variance.

## Acknowledgments

The authors had useful interactions with Robert Schmidt at an early stage of the project. D.A.-G. and A.T. also acknowledge helpful discussions with Alain Barrat, Yehezkel Ben-Ari, Demian Battaglia, Stephen Coombes, Rosa Cossart, Diego Garlaschelli, Stefano Luccioli, Mel MachMahon, Rossana Mastrandea, Ruben Moreno-Bote, and Viktor Jirsa.

## Supporting Information

### TEXT S1

#### Silent Neurons

We define a neuron as silent, if it does not emit more than *S*_Θ_ spikes within the system evolution, which we typically take as the time taken for the network to evolve through to 10^7^ spikes. In particular, in Fig. S 1 (a) and (b) we report the fraction of active neurons *n*\* versus the synaptic strength for two parameter settings and for several values of the considered threshold, namely 0 ≤ *S*_Θ_ ≤ 100. In practice, we observe that neither the minimal value of n* nor the value for which the minimum is reached, appears to strongly depend on the chosen threshold, thus demonstrating the robustness of the results that we present through the article.

#### Mechanisms for the resurgence of silent neurons

In what follows we report the neuronal distributions of the average inter-spike intervals 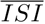, of the corresponding *CV*, of the associated average effective synaptic input 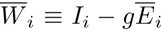 and standard deviation *σ*(*W_i_*). In particular, we consider these distributions for two different stimuli dispersion (namely, Δ*V* = 5 mV and 1mV) as well as for two synaptic strengths (namely, *g* ≃ *g_min_* and *g* >> *g_min_*).

For Δ*V* = 5 mV (Δ*V* = 1 mV) we examine two synaptic strengths, one in proximity of the minimum *g_min_* of *n*\*, where almost 50 % of neurons are active, and one for which almost all the neurons are active again, namely *g* = 4 and 10 (*g* =1 and 4). Let us first consider the distribution of the average 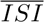 of the single cell reported in Fig S 2(a) and (d). At small *g* the distributions reveal a clear peak at some low 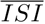 plus a long tail. In correspondence of this coupling the distribution of *CV* is clearly bimodal as shown in Fig S 2(b) and (e) with peaks around zero and one, thus indicating that the neurons associated to the peak in 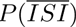 are firing in a regular fashion, while the neurons in the tail of 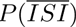 contributes to the second peak in *P*(*CV*) around *CV* ≃ 1. Furthermore, by examining the distributions of the average effective input 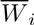 perceived by each single cell, the PDF for *g* ≃ *g_min_* has a peak in proximity the threshold value *V_th_*.

We can conclude that the neurons contributing to the main peaks in 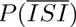 and *P*(*CV*) for *g* ≃ g_min_ are the *winners*, which fire faster than the others and almost periodically, thus suggesting that they are not particularly influenced by the other neurons in the network. Moreover, they correspond to the neurons which are on average above threshold, as shown in Fig. S 2(c) and (f). The neurons contributing to the second maximum in *P*(*CV*) and to the tail of 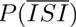 are instead slow neurons whose activity is strongly depressed by the winners and they are neurons around, or just below threshold, in Fig. S 2(c) and (f).

As one can appreciate from Fig. S 2(a) and (d) the 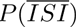 is completely modified at large g. In such a case, a broad peak is present extending over two orders of magnitude. In this regime the majority of the cells are on average below-threshold, as it can be appreciated by the corresponding 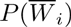, reported in Fig. S 2(c) and (f) as red empty squares, which reveal an almost Gaussian shape centered well below threshold. Therefore we are now in a situation where all neurons are active, but the majority are activated due to the fluctuations in the input and they are no more tonically firing. The fact that now the activity is mostly fluctuation driven, is reflected also in the *CV* distributions, which are now centered well above one.

The reported results clearly show that for wider dispersion of the *I_i_*, as measured by Δ*V*, a greater lateral inhibition is required to observe similar effects.

#### Linear stability analysis

One of the questions that we would like to address is whether the existence of a bursting correlated activity is related to linear stability properties of the network or not. To characterize these properties, we calculate the maximal Lyapunov exponent (LE) λ for the parameters examined in the text. In order to compute the LE we derive from Eq. (3) (main text) its linearization, which describes the evolution of infinitesimal perturbations in the reference orbits, this reads as:

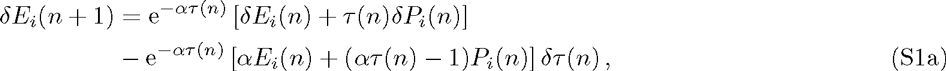

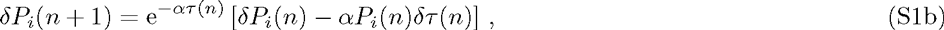

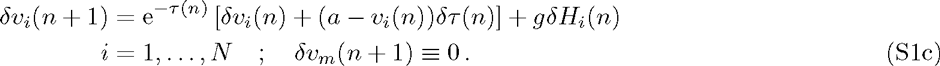

The boundary condition 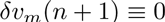 is a consequence of the event driven evolution. The expression of *δτ*(*n*) can be computed by differentiating Eqs. (4) and (5) (in main text), namely

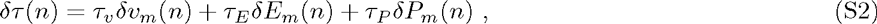

where

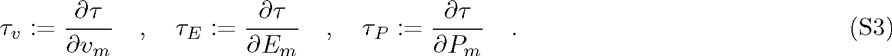

The maximal LE λ is defined as the the average exponential growth rate of the infinitesimal perturbation

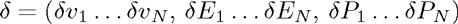

measured through the equation

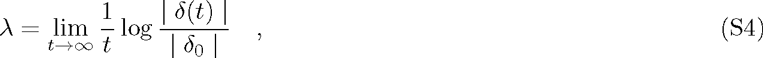

where δ_0_ is the initial perturbation. The evolution of the perturbation δ(*t*) at the following times can be obtained by integrating Eqs. (S1) in the tangent space in parallel with the evolution in the real space and by performing at regular time intervals the rescaling of its amplitude to avoid numerical artifacts, as detailed in [2]. A positive λ denotes a chaotic dynamics, a zero maximal LE is associated to a periodic (or quasiperiodic) orbit, and a negative one to a stable fixed point. It is important to stress that, since we are dealing with an event driven map formulation of the dynamics, the zero Lyapunov exponent which is always present for continuous time evolution and associated to the growth rate of a perturbation along the orbit, is automatically discarded. This implies that, if the evolution is stable, either a fixed point or a periodic solution, we measure in both cases a maximal LE λ < 0.

For a fixed pulse duration *τ_α_* = 20 ms, the behaviour of the maximal LE λ as a function of the coupling *g*, for different excitability spreading Δ*V*, is definitely different. As shown in Fig S3 (a), for Δ*V* = 1 mV the LE (as expected) is zero for very weakly coupled systems, then it first increases with *g* and reaches a maximum around *g* = 2 and then it decreases monotonically becoming negative for *g* > 5. For Δ*V* = 5 mV, the LE is always positive and increases with *g* saturating at an almost constant value λ ≃ 3.4 Hz for *g* ≥ 6. We are specifically interested in the conditions for which the measure *Q*_0_ is maximized, these points are indicated in Fig S3 (a), as one can notice they correspond for both considered Δ*V* to positive λ.

Additionally we have analyzed the behaviour of λ as a function of *τ_α_* by fixing *g* to the value that maximizes *Q*_0_ in the previous analysis. In this case it appears that λ increases with *τ_α_* and becomes definitely negative for sufficiently small *τ_α_* (as shown in Fig. S3 (b)), in agreement with the results reported in [1, 3]. The cell assembly dynamics of our network resembles that of MSNs for large τ_α_, as explained in the text, the point where *Q*_0_ is maximal are indicated also in Fig. S3 (b). These evidences seem to suggest that the striatally relevant dynamics correspond to a chaotic regime, but located in proximity of the transition between chaotic and non-chaotic evolution. The same conclusion was already reported for a rate model of the striatum in [6].

However, all this analysis and the one reported in [6] consider only infinitesimal perturbations, while it has been clearly demonstrated that for inhibitory networks finite perturbations play a fundamental role as shown in [1, 3, 4, 7]. In particular our model, even for λ < 0, can display erratic evolution almost indistinguishable from chaos due to the so-called Stable Chaos mechanism [1, 5]. This leads us to conclude that the usual Lyapunov exponent is unable to capture the degree of erratic motion present in these systems, due to the possible amplification of finite amplitude perturbations.

#### State Transition Matrices for different regimes

In the main text we have just reported the averaged State Transition Matrix (STM) corresponding to the consecutive presentation of two stimuli for parameters obtained by maximizing *Q*_0_. Here we want to show how the STM is modified by considering *τ_α_* = 20 ms, for which *Q*_0_ is maximal, and for a smaller pulse duration, namely *τ_α_* = 2 ms, for which the evolution of the network is seemingly Poissonian. The upper panel of Fig S5 show another realization of the network obtained for the same parameters of Fig. 5 (in main text). The lower panels correspond to *τ_α_* = 2ms. The raster plots clearly show that for *τ_α_* = 20 ms the network exhibits a clear patterned activity with frequent switch from an activated assembly to another, furthermore there is a low correlation between the network activities in presence of the two different stimuli. As shown in Fig S 5 (b,c). For *τ_α_* = 2 ms the system presents much less variability. While it is still capable of discriminating between two different stimuli, now the system fails in revealing a clear assembly switching during the presentation of a single stimulus (see lower panels of Fig. S 5).

#### Synchronized Event Transition Matrices and number of coactive cells for different network realizations

We present two different realizations of the numerical experiment performed in the sub-section *Physiological rele¬vance for biological networks under different experimental conditions.* The difference between the realizations lies on the random connectivity matrix *C_ij_*, which is generated at each realization with the same connection probability. The results are presented in Fig. S 7. More precisely, in Figs. S 7 (a) and S 7 (e) are reported the SETMs for maximal *Q*_0_ (*g* = 8 for the chosen parameters). These are characterized by a large variability in their elements when compared with the corresponding SETMs obtained for decreased inhibition (namely, *g* =1), shown in Figs. S 7 (c) and S 7 (g), is always smaller. The difference between the two regimes is also evidenced in the number of coactive cells: at maximal *Q*_0_ each state is well defined, as illustrated in Figs. S 7 (b) and S 7 (f). Since diagonal elements (representing the number of neurons active in a given state) present larger bars compared with the off-diagonal ones (representing the overlap between two different states) Instead, in the set-up with *g* = 1 the states are hardly distinguishable, diagonal and off-diagonal bars have similar heights (as shown in Figs. S 7d) and S 7h))

**Fig S 1.**
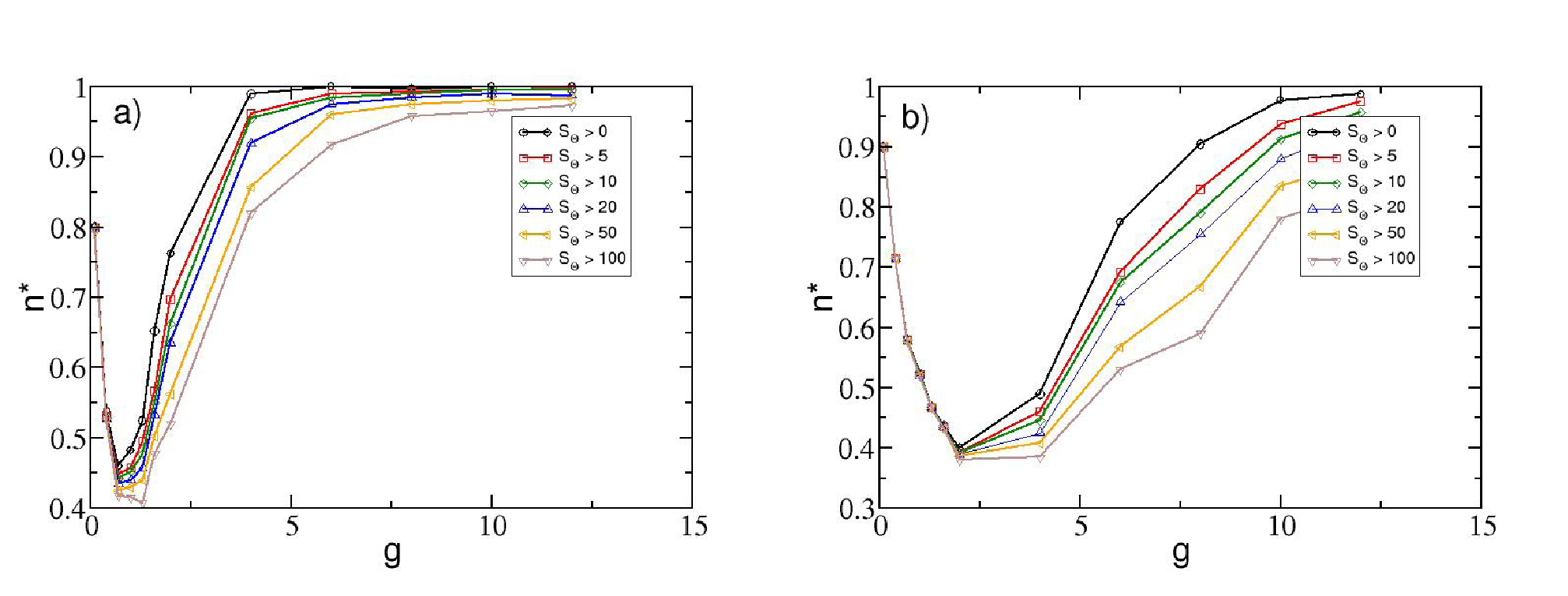
Dependence of the value *n*\* on the chosen threshold. *S*_Θ_ Fraction of active neurons *n*\* vs the synaptic strength, for several threshold definitions. A neuron is considered silent whenever it does not spike at least *S*_Θ_-times during the observation time. Panel a) for Δ*V* = 1 mV and b) for Δ*V* = 5 mV. The system is left to evolve during 10^7^ spikes, after discarding 10^5^ spike events of transient. Other parameters used in the simulation: *K* = 20, *N* = 400 and *τ_α_* = 20ms.

**Fig S 2.**
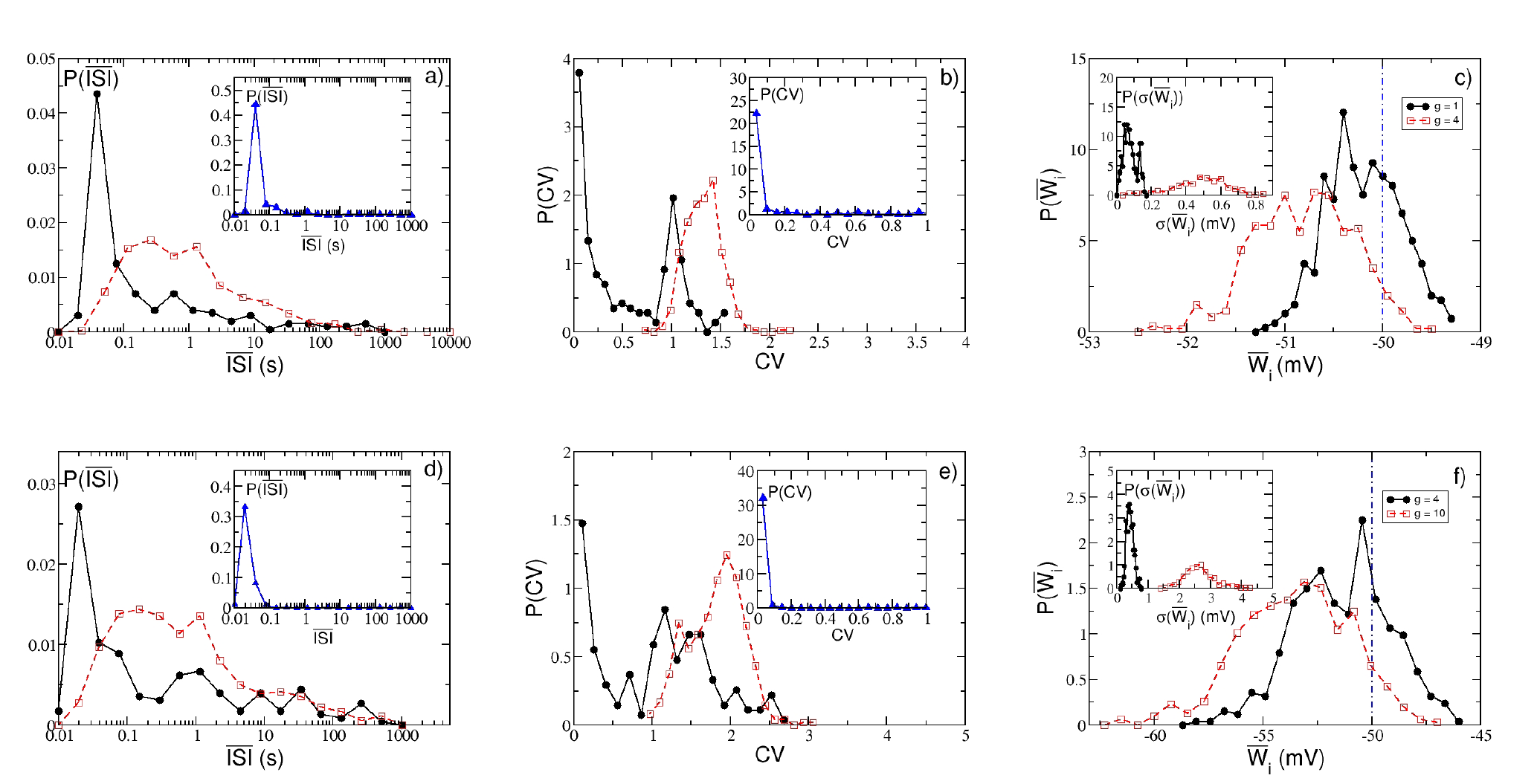
Neuronal statistics. Neuronal distributions of the average 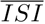 (a,d), of the coefficient of variation *CV* (b,e) and of the average effective synaptic input 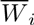 (c,f). The data in the first row are for Δ*V* = 1 mV and in the second one for Δ*V* = 5 mV. For Δ*V* = 1 mV (Δ*V* = 5 mV) black solid line with filled circles correspond to *g* = 1 (*g* = 4) and red dashed lines with open squares to *g* = 4 (*g* = 10). Insets of (a,d) and (c,f), same as the main figure in a lower *g* regime: *g* = 0.4 (*g* = 1) for Δ*V* = 1 mV (Δ*V* = 5 mV). The system is left to evolve during 10^7^ spikes, after discarding 10^5^ spike events. Other parameters used in the simulation: *K* = 20, *N* = 400 and *τ_α_* = 20 ms.

**Fig S 3.**
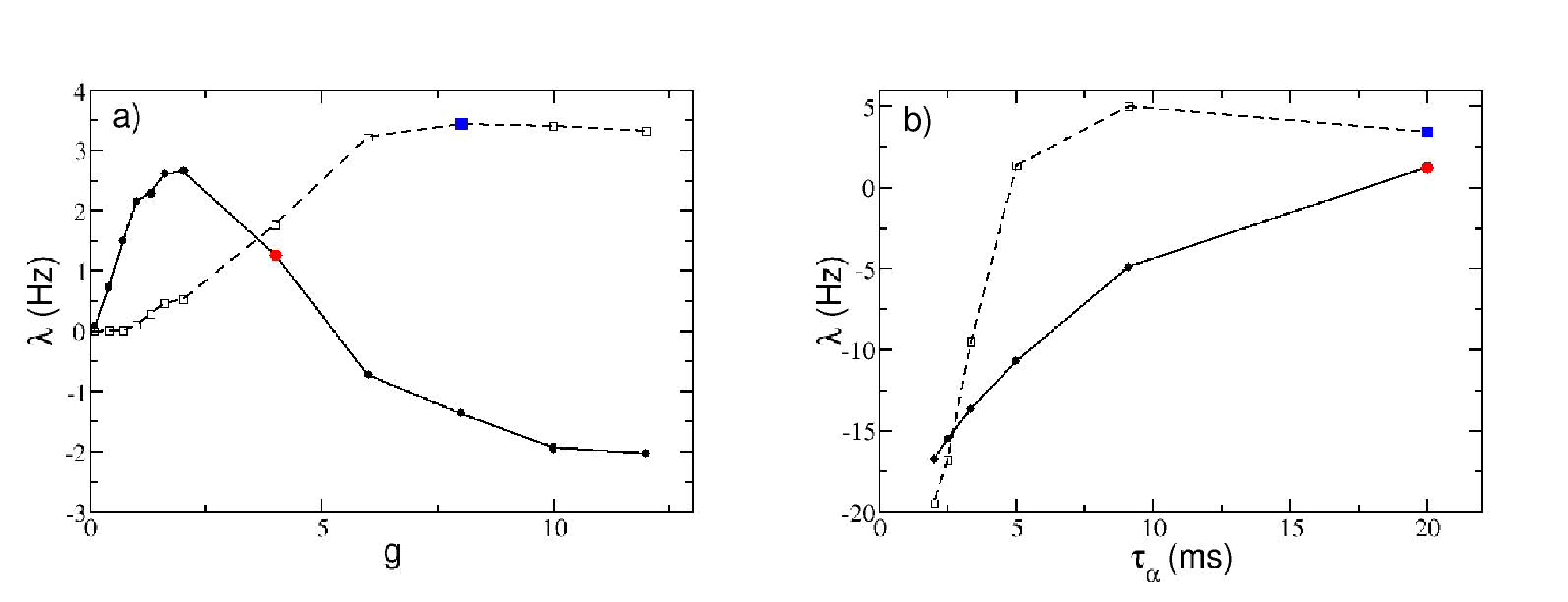
Linear stability analysis. a) Maximal Lyapunov exponent λ at a fixed *τ_α_* = 20, as a function of the synaptic strength for Δ*V* = 1 mV (continuous line, filled circles) and Δ*V* = 5 mV (dashed line, empty squares). b) Maximal Lyapunov exponent λ as a function of the pulse duration *τ_α_* for the parameters 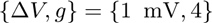 (continuous line with filled circles) and {5 mV, 8} (dashed line with empty squares). In both panels, the blue filled square indicates the triad 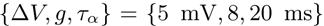, and the red filled circle to 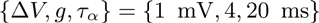; these values are associated to the maximum values of *Q*_0_ obtained for excitability distributions with fixed width Δ*V*. The tangent space Eq. (S1) is evolved during a period corresponding to 10^6^ spikes, after discarding a transient of 10^5^ spikes. Other parameters used in the simulation: *K* = 20, *N* = 400.

**Fig S 4.**
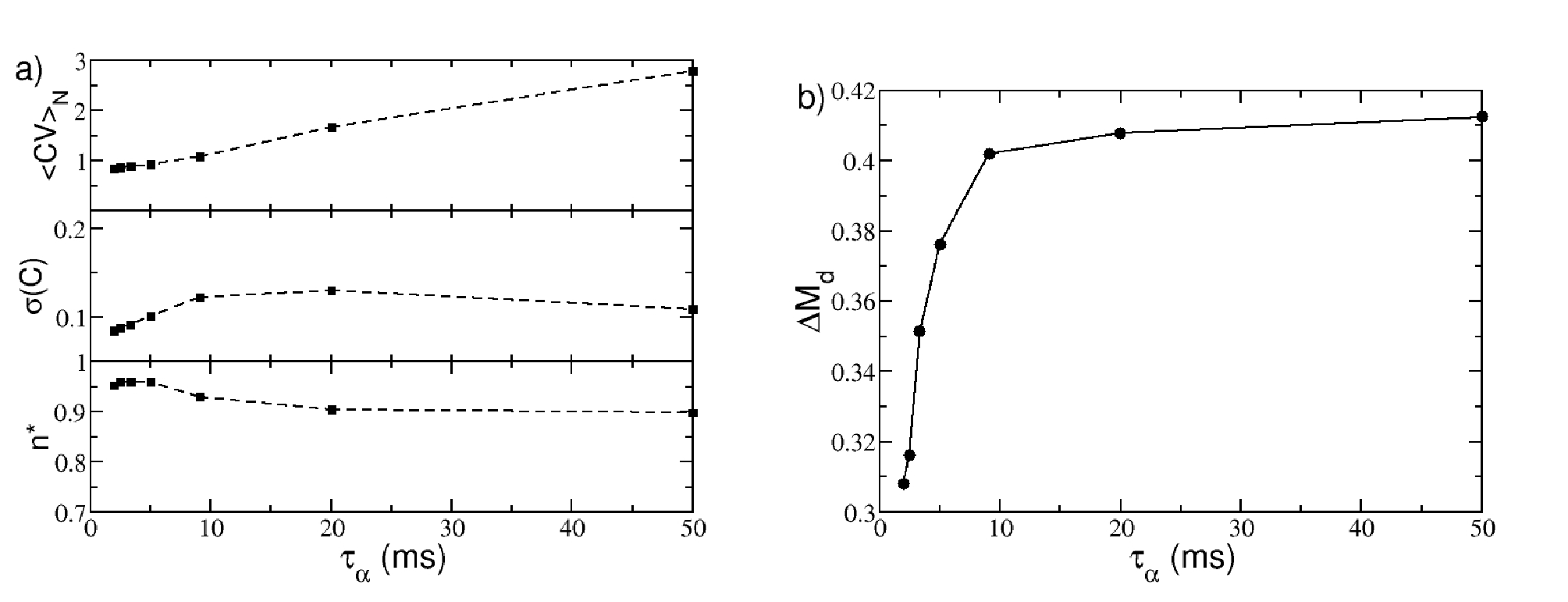
Metrics of the structured activity vs synaptic time decay. a) Metrics entering in the definition of *Q*_0_ and their dependence from τ_α_. From top to bottom: Averaged coefficient of variation ⟨*CV*⟩_*N*_, standard deviation of the cross-correlation matrix *σ*(*C*), and the fraction of active neurons *n*\*. b) Δ*M_d_* as a function of *τ_α_*. The system is left to evolve during 10^7^ spikes, after discarding 10^5^ transient spike events. Parameters here used Δ*V* = 5 mV, *g* = 8, *K* = 20, *N* = 400.

**Fig S 5.**
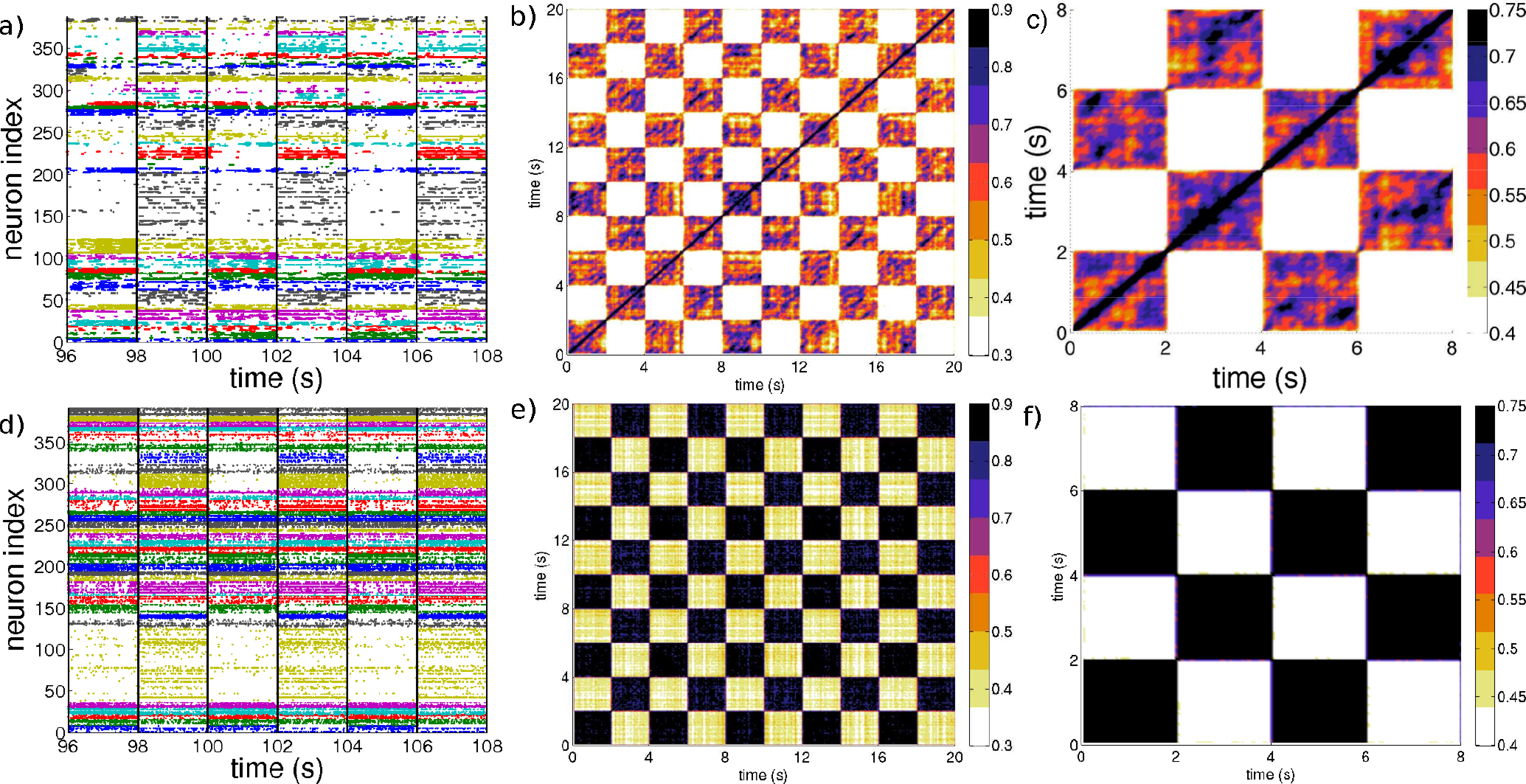
Two stimuli presentation. Upper panel, another realization of the network with the same parameters as chosen in the main text and *τ_α_* = 20ms. Lower panel, network with *τ_α_* = 2ms. From left to right it is depicted the raster plot colored according to the *k-means* algorithm with *k*=25, vertical lines indicates the change of the presented stimulus. In the middle column is reported the state transition matrix calculated over a time span of 20 seconds (the stimulation protocol is repeated 5 times). Rightmost column reports the state transition matrix for a block 4 s χ 4 s averaged over *r* = 5 successive presentations of the inputs.

**Fig S 6.**
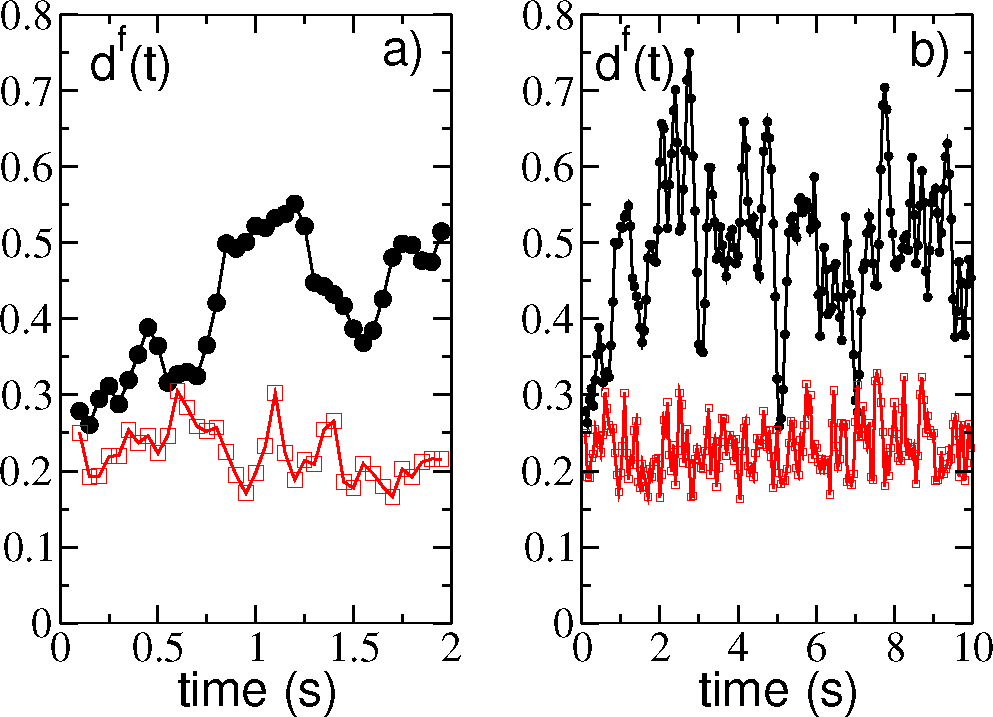
Pattern Separation. Dissimilarity measure in time for an observation window of length a) *T_E_* = 2 s and b) *T_E_* = 10 s, for two values of *τ_α_* = 20 ms (black circles) and *τ_α_* = 2 ms (red squares) at a fixed value of *f* = 0.2. It is clearly observed that *τ_α_* = 20 ms more effectively differentiates the similar inputs in both observation windows, as seen by the larger values of dissimilarity respect to the *τ_α_* = 2ms. The initial increase of *d^f^*(*t*) observable for *τ_α_* = 20 ms in panel (a) is probably due to the fact that the dynamics for this choice of parameters is chaotic as shown in the *Linear stability analysis* sub-section. Therefore the increase can be associated to a transient evolution towards the final attractor. Other parameters used: Δ*T* = 50 ms, *g* = 8, *N* = 400, *K* = 20.

**Fig S 7.**
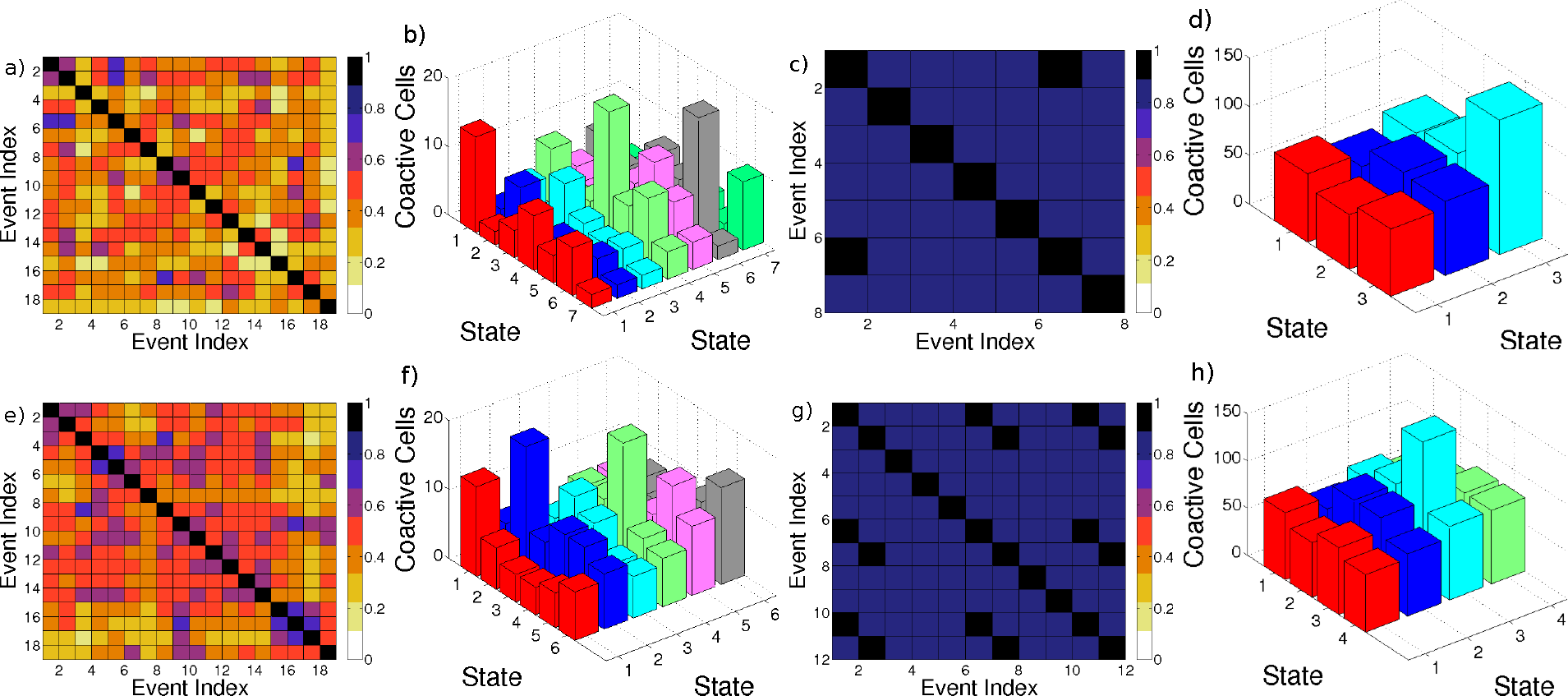
Response of the network to an increase in the excitability. Two different realizations of the numerical experiment reported in the subsection *Physiological relevance for biological networks under different experimental conditions* of the main text. First (third) column reports two realizations of the SETM estimated for *g* = 8 (*g* = 1). Second (fourth) column displays the number of coactive cells in the corresponding cases for *g* = 8 (*g* = 1). The other parameters for the reported simulations are Δ*V* = 5 mV, *K* = 20, *N* = 400.

